# Phenotypic similarity of NAD(P)-Malic Enzymes in Tomato: Unveiling Patterns of Convergent and Parallel Evolution

**DOI:** 10.1101/2025.02.12.637823

**Authors:** Andrea Martinatto, Jonas M. Böhm, Claudia A. Bustamante, Francesco Pancaldi, Eric Schranz, Marcos A. Tronconi

**Author notes:** These authors have contributed equally to this work (Andrea Martinatto, Jonas M. Böhm).

## Abstract

The *Solanum lycopersicum* (tomato) genome encodes seven malic enzymes (MEs): three cytosolic NADP-ME isoforms (SlNADP-ME1, -2, and -3), two plastidic NADP-ME proteins (SlNADP-ME4a and -4b), and two mitochondrial NAD-dependent subunits (SlNAD-ME1 and -2) of a heteromeric enzyme. Except for SlNADP-ME3, which is almost exclusively expressed in the root, all the genes are active in the major tissues of flowering plants (leaf, stem, flower, and root), with SlNADP-ME2 and -4b transcripts accumulating at relative high levels. During the cell expansion and ripening phases of fruit development, *SlNADP-ME3* and *-4b* expression increased in the pericarp and seeds while *SlNADP-ME2* remained at a high level. In green fruit, *SlNADP-ME3* and *-4b* are down-regulated by ethylene, with *SlNAD-ME1* and *-2* being up-regulated. A correlation between SlNADP-ME4b accumulation and NADP-ME activity was observed from the early immature green to the mature green fruit stages, linking this plastidic isoform to starch and lipid biosynthesis. *SlNADP-ME1* and *-4a* expression increase with temperature, suggesting involvement in defense mechanisms and supported by *cis*-elements composition in their promoters. Interesting, SlNADP-ME3 biochemical properties and its accumulation in seeds is part of an inherited and long-conserved genetic lineage in Angiosperms including Arabidopsis NADP-ME1, involved in normal seed germination. By analyzing the phylogeny and synteny of *SlNAD(P)-ME* genes and determining the biochemical properties of the recombinant proteins, we seek functional similarities to Arabidopsis NAD(P)-ME members. Our findings enhance the understanding of malate metabolism in tomatoes, which could inform strategies to improve fruit quality. Additionally, they prompt a re-evaluation of pre-existing notions regarding the functional orthology of enzymes based on phylogenetic relationships.

**SIGNIFICANCE STATEMENT:** Enzyme functional orthology among distant plant species cannot be inferred through phylogenetic analysis because of the large independent rearrangements of genomes during evolution. A comprehensive characterization of the tomato NAD(P)-ME family and their comparison with members from an unrelated plant species, such as Arabidopsis, allowed us to evaluate the impact of selective pressures on orthologous genes. This approach reveals that the phenotypic similarity of an entire enzyme family in different species is achieved through diverse evolutionary events such as ancestry, parallelism, and genetic convergence.

## INTRODUCTION

Tomato (*Solanum lycopersicum* L.) is the second world’s leading fruit (or vegetable) crop next to potato (*Solanum tuberosum*). Tomato fruit is a fundamental nutritional component of a balanced human diet (Martí et al., 2016) due to its numerous health-promoting compounds (vitamins, carotenoids and phenolic compounds). These bioactive compounds have a wide range of physiological properties, including antioxidant, anti-allergic, antimicrobial, anti-inflammatory, vasodilator and cardioprotective effects (Raiola et al., 2014). Besides its economic and nutritional importance, tomato is the model plant for fleshy fruit development studies.

After flower fertilization, fruit development begins with an extraordinary metamorphosis of the ovary that includes the enlargement of the placenta and the pericarp, the outermost tissue of the fruit that defines the locular cavity where all other tissues reside (Giovannoni, 2004; Tanksley, 2004; Matas et al., 2011). During pericarp development, the cells of the epidermis and adjacent exocarp divide generating additional cell layers through lipid biosynthesis and membrane deposition allowing fruit growth. Meanwhile, the cells of the mesocarp enlarge, mainly due to a massive influx of water into the vacuole, and endocarp (or inner epidermis) remains as a single cell layer (Matas et al.,2011). Starch, sucrose, hexoses, organic and amino acids are accumulated in mesocarp derived from the photosynthetic activity of exocarp (Matas et al., 2011). Malate is the second most abundant organic acid after citrate; during the expansion phase of tomato fruit, malate contributes significantly to the osmotic potential driving cell enlargement through water uptake (Liu et al., 2007; Biais et al., 2014). Also, along with citrate and hexoses, malate determines the total soluble solids content, a key factor in fruit taste and processing quality (Quinet et al., 2019; Fridman et al., 2000). The accumulation pattern of malate in tomato fruits exhibits a biphasic profile, with maximum levels reached during the cell expansion phase followed by a decrease during ripening (Carrari et al., 2006; Oms-Oliu et al., 2011). This dynamic suggests this acid may serve as a precursor for the biosynthesis of other metabolites accumulating during the last phase of fruit maturation, such as hexoses, citrate, aspartate or glutamate (Biais et al., 2014). In addition, malate could be oxidized through the alternative mitochondrial respiratory chain (Wedding, 1989) to meet the high ripening demands of climacteric fruit. Malate metabolism in tomato would be ethylene-driven, as its accumulation decreases in LeACS2 antisense plants, impaired in ethylene biosynthesis, when exposed to ethylene (Gao et al., 2007). The characteristic red tomato color is a result of the accumulation of the carotenoid lycopene in the epidermis, exo- and mesocarp. Flavonoids, alkaloids, phenolic and volatiles compounds biosynthesis is restricted to epidermis and exocarp, while the deposition of a cuticle layer on the fruit surface is a functional specialization of the epidermis (Mintz-Oron et al., 2008; Adato et al., 2009; Quinet et al., 2019). During ripening, the mesocarp constitutes the metabolic factory of the pericarp, with genes encoding transporters and enzymes for starch, sucrose and TCA cycle intermediates being up-regulated; at the time that the cell walls undergo changes that result in fruit softening (Vicente et al., 2007; Saladié et al., 2007). Finally, the endocarp produces an inner cuticle layer that restricts the movement of metabolites, gases, or fluids between the pericarp and the locular cavity, defining the biomechanical properties of ripe fruit (Matas et al.,2011; Tamasi et al., 2016; Quinet et al., 2019).

In plants, key enzymes involved in malate oxidation include NADP-dependent malic enzyme (NADP-ME) and NAD-dependent malic enzyme (NAD-ME), which catalyze the oxidative decarboxylation of malate to produce pyruvate, CO2, and NAD(P)H (Drincovich et al.,2001; Maurino et al., 2009). NADP-ME is found in cytosol and plastid while NAD-ME is present in mitochondria (Lai et al., 2002a, 2002b; Gerrard Wheeler et al., 2005; Tronconi et al., 2018). Plant NADP-ME is assembled as homooligomers (dimers and tetramers) of monomers of ∼62-65 kDa with several isoforms distributed over plant tissues (Chi et al., 2004; Gerrard Wheeler et al., 2005; Maurino et al., 2009; Alvarez at el., 2013; Saigo at al., 2013). On the other hand, mitochondrial NAD-ME (mNAD-ME) is a heteromer of two distinct proteins (∼65% sequence identity): the α-subunit (or NAD-ME1) and β-subunit (or NAD-ME2) with molecular masses of 63 and 58 kDa, respectively (Willeford and Wedding, 1987; Tronconi at al., 2008, 2010a, 2018). mNAD-ME is ubiquitously present in photosynthetic and non-photosynthetic cells of higher plants (Tronconi et al., 2010b; Hüdig et al., 2022).

The eudicot *Arabidopsis thaliana* possesses four *NADP-ME* genes encoding three cytosolic NADP-MEs (*AtNADP-ME1*, *-2* and *-3*) and one plastidic isoform (*AtNADP-ME4*). *AtNADP-ME2* and -*4* genes are constitutively expressed in vegetative and reproductive organs, whereas the expression of At*NADP-ME1* and At*NADP-ME3* is regulated by both developmental and cell specific signals (Gerrard Wheeler et at., 2005). Specifically, AtNADP-ME2 is responsible for the bulk of NADP-ME activity in leaves playing a role during the basal defense response, where it is required to produce ROS following pathogen recognition (Voll. et al. 2012). The expression of *AtNADP-ME3* is restricted to trichomes and pollen, while *AtNADP-ME1* is expressed in the embryo at the late stages of embryogenesis and seed maturation, and in the root tip during germination (Yazdanpanah et al., 2018; Gerrard Wheeler et at., 2005). *AtNADP-ME1* loss-of-function mutant plants display reduced seed viability and germination (Arias et al., 2018; Yazdanpanah et al., 2018) and enhanced tolerance to Al^+3^ stress (Badia et al., 2020), highlighting the hierarchy of NADP-ME activity in the physiological background in which is expressed. The recombinant AtNADP-ME proteins exhibit different kinetic properties and metabolite regulation (Garrard Wheeler et al., 2008; Maurino et al., 2009). Considering this, the different NADP-ME isoforms from *A. thaliana* can be assigned specific rather than redundant functions (Maurino et al., 2009). Mitochondrial NAD-ME is the most versatile member of the plant NAD(P)-ME family, mostly structured as an αβNAD-ME heteromer but also as α2 and β2 homomers, which have been detected in specific Arabidopsis tissues (Tronconi et al., 2010a). Each NAD-ME entity differs in its kinetic mechanisms and regulation by metabolic effectors (Tronconi et al., 2008, 2010b, 2015) to supply pyruvate during citrate and amino acid production (Sweetlove et al., 2010; Lehmann et al., 2016; Le et al., 2022) or drive malate-dependent alternative respiration activity (Sweetman et al., 2009).

Despite the wealth of information available on malic enzymes in *A. thaliana*, our understanding of malate metabolism in fruit remains limited by the lack of information on the biochemical properties of tomato NAD(P)-ME isoforms. *S. lycopersicum* and *A. thaliana* belong to the Solanaceae and Brassicaceae families, respectively, which diverged early in the eudicot radiation (∼130 million years ago) (Ku et al., 2000). The evolutionary distance between both families, coupled with differences in genome evolution and selective pressures, makes it challenging to infer functional orthology based solely on sequence homology (Fridman and Zamir, 2003). The evolutionary dynamics of the two genomes have been traced by different molecular events and selective pressures. In Brassicaceae, two rounds of whole genome duplication (WGD), chromosomal rearrangements and selective loss of extra copies of duplicated genes were the main processes leading to rapid speciation in the family (Barker et al., 2009). In contrast, the Solanaceae genome diversified in the absence of WGD and evolved through small-scale genomic events (Wang et al. 2018). Because homoplasy (parallel and convergent evolution) is a common feature when similar evolutionary constraints operate over different genetic backgrounds (Fridman and Zamir, 2003), functional orthology cannot be directly assigned to proteins by analysis of phylogenetic relationships. Thus, the idea of “phenotypic similarity” emerges as a concept that defines the evolution of similar adaptive solutions because of environmental challenges, independently of phylogenetic relationships (Cerca, 2023). This study aims to gain insights into distinctives properties of NAD(P)-ME isoforms in tomato plants (SlNAD(P)-ME), mainly through analyses during fruit development and ripening. We evaluated changes in enzyme activity, protein abundance, and transcript levels of SlNAD(P)-ME isoforms during pericarp development and the effect of ethylene inhibition to identify tissue-specific isoforms and their potential functions. By analyzing the phylogeny, synteny, and promoter regions of *SlNAD(P)-ME* genes, and determining the biochemical properties of the recombinant proteins, we seek functional similarities to Arabidopsis *NAD(P)-ME* members. Our findings contribute to a more comprehensive understanding of malate metabolism in tomato, potentially informing strategies to improve fruit quality and postharvest characteristics, but also important takes a look at pre-established ideas about the functional orthology of enzymes based on phylogeny.

## RESULTS

### Developmental changes in NAD(P)-ME activities in tomato fruits

We measured NADP-ME and NAD-ME activities in tomato fruit pericarp at seven stages of development spanning the phases of cell expansion and ripening (i.e. Early Immature Green [EIG, 10 DPA], Late Immature Green [LIG, 30 DPA], Mature Green [MG, 38 DPA], Turning [T, 42 DPA], Orange [O, 46 DPA], Red Ripe [RR, 48 DPA] and Red Ripe after 7days [RR+ 7, 55 DPA ]) (Figure 1A). When NADP-ME and NAD-ME activities are expressed on a fresh weight (FW) basis, both showed a similar overall trend; the activities are highest at 10 DPA and then decrease during the following stages (Figure 1B). At the end of fruit development (RR +7), the FW-based NADP-ME and NAD-ME activities were 5.5 and 2-fold lower than at the very early stage of development (EIG), respectively (Figure 1B).

**Figure 1.**
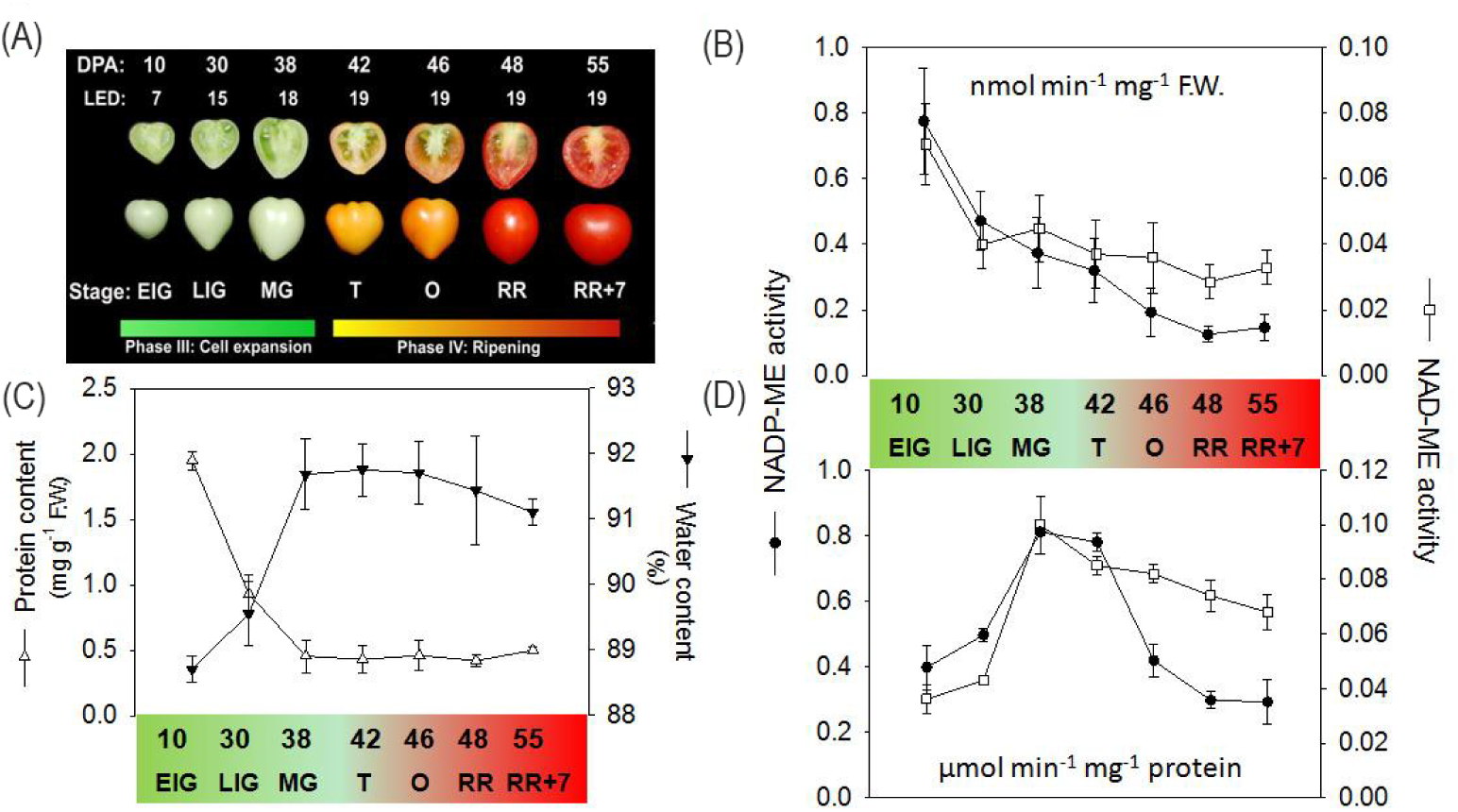
NADP-ME and NAD-ME activities during tomato pericarp development. (**A**) Development stages of tomato fruit from Micro-Tom cultivar analyzed in this work. Fruits were harvested at the indicated days after anthesis (DPA). The larger equatorial diameter (LED, mm) of fruit is depicted. (**B**) and (**D**) Maximal activities were measured in fruit pericarps harvested between 10 - 55 DPA and expressed as nanomoles per minute per gram fresh weight (**B**) or micromoles per minute per microgram of protein (**D**). (**C**) Water percentage and total protein content. Supplementary Data represents the mean ± s.d (standard deviation) of three to five independent measurements.

The increase in water content and the drastic decrease in total protein during fruit development (Figure 1C) shape the enzymatic activity profiles (Biais et al., 2014) and likely obscure any stage-specific role of NAD(P)-ME. Indeed, when the enzymatic activities are expressed on a protein basis, a different pattern of activities was observed for NADP-ME and NAD-ME (Figure 1D). NADP-ME activity was low in the early stages (10-30 DPA), then increased and reached a plateau between the MG and T stages. The enzymatic activity then dropped significantly during the maturation phase. On the other hand, the NAD-ME activity remained very low until 30 DPA followed by an abrupt 3-fold increase at the MG stage. Thereafter, NAD-ME activity decreased slowly but gradually until 55 DPA. Thus, the NADP-ME activity at the RR+7 stage reached levels that were similar to those at the EIG stage (10 DPA), whereas the NAD-ME activity remained at least twice as high when comparing the same stages (Figure 1D).

We explored the presence of NAD(P)-ME isoforms at different stages of pericarp development using native and/or denaturing (SDS) PAGEs. The abundance of total immunoreactive tomato NADP-ME proteins detected by western blot after SDS-PAGE showed no significant changes during pericarp development (Figure 2A). As NADP-ME entities can exhibit different electrophoretic mobilities (Gerrad Wheeler et al., 2005), we analysed NADP-ME activity after native PAGE and found two protein bands of different mobilities (Figure 2B). A low-mobility protein band is present in all stages of pericarp development examined and in the tomato leaf (Figure 2B). From 30 DPA up to 55 DPA, an additional band with slightly higher mobility was present. The mobility of both protein bands found in tomato tissues is similar to that of the homotetrameric NADP-ME from maize leaf (Figure 2A), so we assume that they correspond to at least two different tetrameric NADP-ME isoforms expressed in the pericarp. No active bands consistent with dimeric assembly of NADP-ME were observed in the samples.

**Figure 2.**
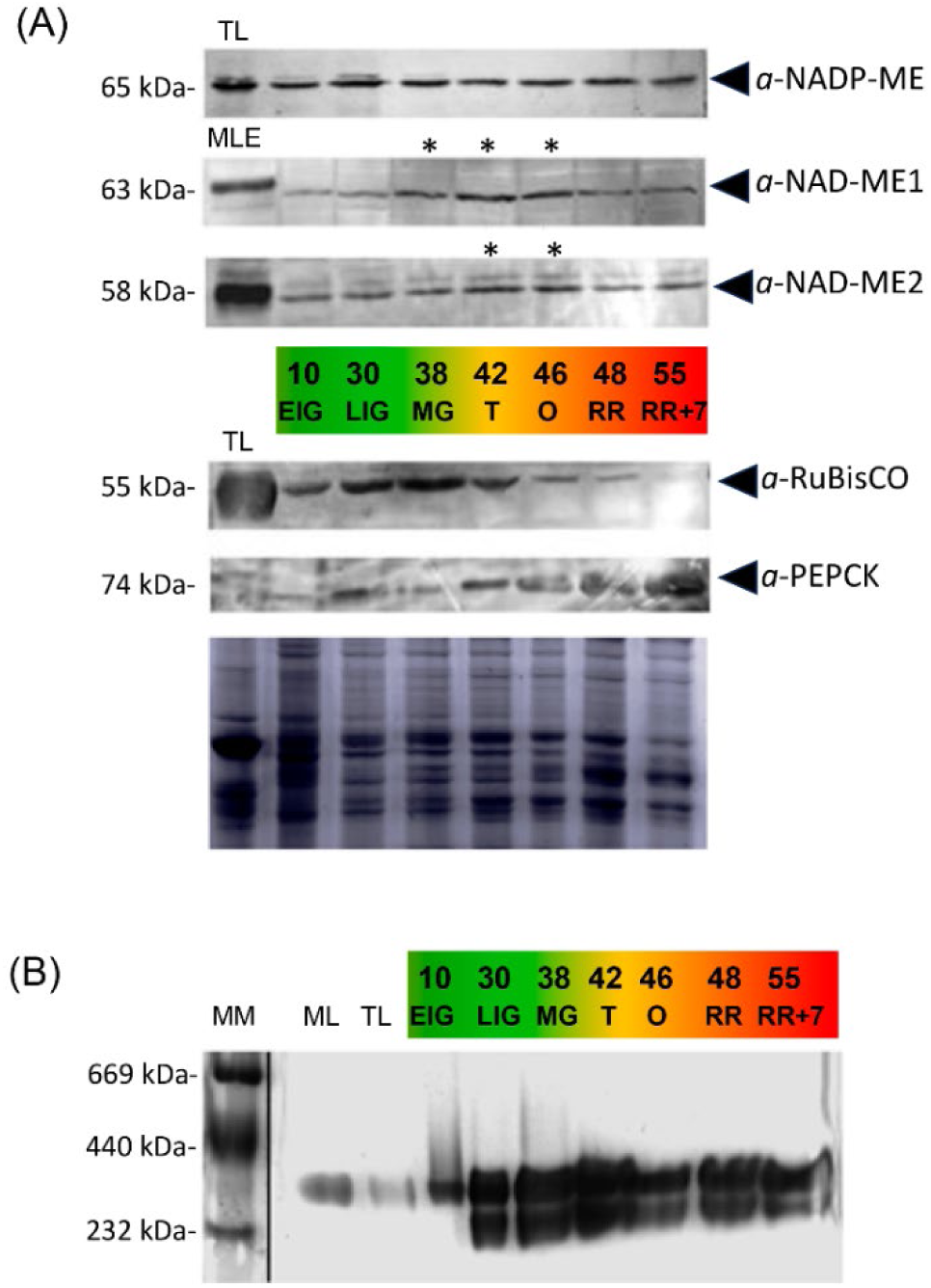
NADP-ME and NAD-ME proteins of tomato fruits pericarp analyzed by PAGE. (**A**) S.DS-PAGE revealed by western blots were conducted using antibodies against NADP-ME, AtNAD-ME1, AtNAD-ME2, RuBisCO and PEPCK. Thirty micrograms of total soluble protein were added per lane. As a control, the same amount of tomato leaf crude extract (TL) was loaded in the first lane for western blot revealed with antibodies NADP-ME, RuBisCO and PEPCK. In the case of western blot revealed with antibodies against AtNAD-ME1 and -2, thirty micrograms of Arabidopsis mitochondrial leaf extracts (MLE) were loaded in the first lane. The molecular masses (kDa) of the immunoreactive bands are shown on the left. Significant differences in the level of the immunoreactive band, assessed by densitometric analysis of at three gels, in relation to the amount in fruits at 10 DPA, are indicated (*) (Mann-Whitney test, p<0.1, n=3). The content of RuBisCO and PEPCK proteins were also analyzed (see Materials and methods). (**B**) Native-PAGE revealed by *in-gel* activity assays. Eight micrograms of total soluble protein from pericarps or tomato leaf (TL) were loaded in each case. In the lane indicated as ML, 4 milliunits of NADP-ME from maize leaf crude extract was loaded. MM: Molecular mass marker.

We used polyclonal antibodies generated against the NAD-ME1 that do not cross react with the NAD-ME2 and vice versa (Tronconi et al., 2008). Therefore, an unequivocal assignment of the immunoreactive protein for mitochondrial isoforms could be established during fruit development. NAD-ME1 and -2 protein levels increased significantly during the cell expansion phase, with a maximum reached between the MG and O stages and a decrease during the maturation phase (Figure 2A). The content of RuBisCO and PEPCK proteins were also analyzed. These enzymes show an opposite accumulation profile, consistent with the re-routing of the carbon flow from photosynthesis to the gluconeogenic pathway during tomato fruit development.

Overall, low NAD(P)-ME activities were observed during the early stages of cell expansion, increasing and reaching a plateau between the MG and O stages, followed by a slight (NAD-ME) or significant (NADP-ME) decrease during ripening.

### Sequence retrieved and phylogenetic relationships of *SLNAD(P)-ME* in eudicots

We retrieved five *NADP-ME* and two *NAD-ME* gene sequences from the tomato genome (Annotation ITAG4.0; http://solgenomics.net) and named them as *SlNADP-ME1* (Solyc08g066360), *SlNADP-ME2* (Solyc05g050120), *SlNADP-ME3* (Solyc12g008430), *SlNADP-ME4a* (Solyc03g120990), *SlNADP-ME4b* (Solyc12g044600), *SlNAD-ME1* (Solyc08g013860) and *SlNAD-ME2* (Solyc01g094200). *SlNADP-ME1*, *-2* and *-3* encode cytosolic isoforms and were named following the criteria for designating their counterparts in *A. thaliana* (Gerrard Wheeler et al., 2005). In this sense, *SlNADP-ME2* and *SlNADP-ME3* encode for proteins closely related (90 % sequence identity) and *SlNADP-ME1* express a protein with the lowest amino acid identity value among cytosolic isoforms (∼78% sequence identity). On the other hand, different prediction programs (ARAMEMNON, http://aramemnon.botanik.uni-koeln.de) indicated that *SlNADP-ME4a* and *-4b* encode plastid-localized isoforms with 87% sequence identity. *SlNAD-ME1* and *-2* genes encode the α- and β-NAD-ME subunits (65% sequence identity), respectively, of the mitochondrial NAD-MEα/β heteromer (Tronconi et al., 2008; Tronconi et al., 2020).

The evolutionary relationship of tomato NADP-ME was addressed using Maximum likelihood (ML, Figure 3) and Bayesian Inference (BI, Figure S1). According to the protein-based ML tree, three main NADP-ME lineages are present in the eudicot’s genome (Figure 3). Lineages II and III constitute sister monophyletic groups of NADP-MEs and Lineage I is defined as a paraphyletic clade grouping ancestral isoforms. The cytosolic NADP-MEs are included in Lineage I and/or II, and Lineage III contains all plastidic NADP-MEs (Figure 3). From this topology, SlNADP-ME1 groups with AtNADP-ME2 and -3 in Clade II, while SlNADP-ME2 and - 3 are part of the ancestral Lineage I that also contains AtNADP-ME1 on a different branch of this paraphyletic clade (Figure 3). In accordance with the above predictions, SlNADP-ME4a and -4b are included in Lineage III of plastidic isoforms.

**Figure 3.**
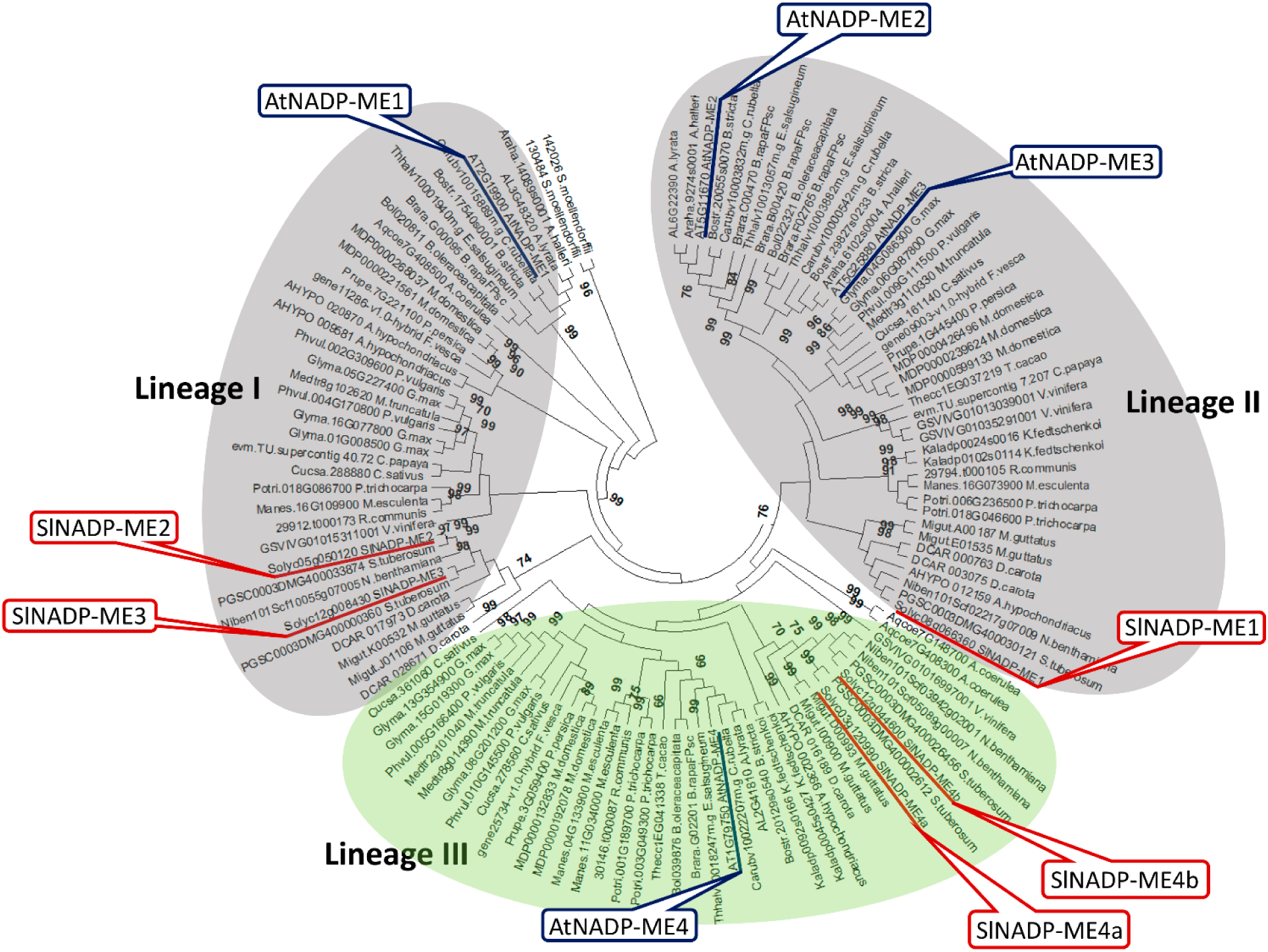
Phylogenetic gene tree of NADP-ME proteins of Eudicotydeloneae. The evolutionary history was inferred using the ML method using a MSA of 347 protein sequences with 546 amino acid positions from genomes from 152 eudicot species and *S. moellendorffii*, an early branching tracheophyte used as outgroup. The NJ optimal tree with the sum of branch length = 1.58 is shown. The tree is drawn to scale, with branch lengths measured in the number of substitutions per site. The evolutionary distances were computed using the JTT matrix-based method with a discrete Gamma distribution to model evolutionary rate differences among sites (G, parameter = 0.52). Support values of 2,000 bootstrap replicates are given as MLB next to the branches when greater than 50%.

To assess whether selective evolution shaped the grouping in the phylogenetic tree inferred from amino acidic NADP-ME sequence, we performed tree constructions using third codon positions, less affected by selection. The BI tree (Figure S1) was overall congruent with the inferred from amino acid sequences except for Lineage I that appears as a monophyletic group. In the IB analysis we globally recovered the expected topology based on published eudicot species phylogenies (Hilu et al., 2014) suggesting that (i) three *NADP-ME* gene linages are present in eudicot genomes, and (ii) some driving force modified the amino acid sequences alternating the relationship between members of Lineage I in the protein-based tree. From these analyses, we suggest that (i) *SlNADP-ME1* is phylogenetically related to *AtNADP-ME2* and *-3* paralogous genes, (ii) *SlNADP-ME3* and *AtNADP-ME1* share a common ancient ancestry and (iii) *SlNADP-ME2* is a neo functionalized gene copy of *SlNADP-ME3*.

In higher plants, *NAD-ME* genes constitute two lineages of paralogs, *α-NAD-ME* and *β-NAD-ME*, phylogenetically unrelated to *NADP-ME* counterparts (Tronconi et al., 2018, 2020). We previously showed that *SlNAD-ME1* and *-2* are included in *α-NAD-ME* and *β-NAD-ME* clades, respectively (Tronconi et al., 2018).

### Synteny Analysis of *NADP-ME* genes in Angiosperms

To further elucidate the evolutionary dynamics and duplication histories of the *NADP-ME* genes of *S. lycopersicum* and *A. thaliana*, the syntenic relationships of the three tomato and the four Arabidopsis *NADP-ME* genes, along with their homologs across 310 angiosperm genomes, were detailly studied (Supplementary Data 1). The results of these analyses showed that the *NADP-ME* genes are overall highly conserved in their genomic positioning across angiosperms. Still, some relevant, lineage-specific, genomic patterns were also unveiled. In total, eight NADP-ME syntenic communities were found across angiosperms (Figure S2 and Supplementary Data 1). Each of these syntenic communities represents an independent “genomic configuration” of a group of *NADP-ME* genes, across specific taxonomic group(s). Out of these 8 syntenic communities, community 1 is conserved across the vast majority of angiosperm species, and groups up to 33% of the species analyzed, including *SlNADP-ME1* and *AtNADP-*ME4 (Supplementary Data 1). Communities 3, 4, and 5 also represent highly conserved genomic configurations of *NADP-ME* genes, overall grouping 61% of the total *NADP-ME* angiosperm copies. However, their conservation is restricted to either monocot (community 5) or dicot species (communities 3 and 4). In addition, community 3 contains *SlNADP-ME3* and *AtNADP-ME2* genes, while community 4 includes *AtNADP-ME1*. Finally, communities 2, 6, 7, and 8 represent more lineage-specific genomic configurations, conserved within the Brassicaceae (communities 2 and 6) and Solanaceae families (community 7), as well as the Papaver genus (community 8). Remarkably, the Solanaceae-specific community 7 contains the *SlNADP-ME2* gene (Figure S2 and Supplementary Data 1). As such, synteny analysis overall shows that while *SlNADP-ME1* and -*3* genes – along with *AtNADP-ME 1 and 2* copies – lie in highly conserved genomic configurations of the *NADP-ME* genes, the *SlNADP-ME2* and *AtNADP-ME3* gene evolved instead toward distinct genomic contexts specific to Solanaceae and Brassicaceae, respectively (Supplementary Data 1). This likely depends on specific chromosomal rearrangements that took place during the evolutionary history of these families and might denote specific functional properties associated with these two gene copies.

### Expression profiles of *SlNAD(P)-ME* genes during tomato fruit development

All *SINAD(P)-ME* genes are transcribed, although at different levels (Figure S3). SlNADP-ME2 and SlNADP-ME4b transcripts accumulate at high levels in leaves, stems, flowers, and roots (Figure S3). SlNADP-ME1 and -4a are also present in these organs but at much lower levels. SlNADP-ME3 transcripts are absent in flowers, while transcript abundance is high in roots and very low in leaves and stems (Figure S3). *SlNAD-ME1* and *-2* are constitutively expressed at similar levels in photosynthetic and non-photosynthetic tissues, consistent with the housekeeping role of heteromeric NAD-MEα/β in plant mitochondria (Tronconi et al., 2008; Le at al., 2022).

To identify the specific NAD(P)-ME isoforms that could generate the observed activity profiles, we analyzed the pattern of expression of all *NAD(P)-ME* genes at the seven stages of pericarp fruit development (Figure 4). We found that *SlNADP-ME2* and *SlNADP-ME4b* are the most highly expressed genes (1.4 to 10-fold higher than *UBQ3*). We also found that, except for *SlNADP-ME2*, which expression showed the lowest variation during fruit development, the changes in the expression pattern of the other *SlNAD(P)-ME* genes could be assigned to the two developmental phases assayed, cell expansion and ripening. *SlNADP-ME1*, *SlNADP-ME4a*, *SlNAD-ME1* and *SlNAD-ME2* have a higher expression during the cell expansion phase than that during ripening (Figure 4). In contrast*, SlNADP-ME3* and *SlNADP-ME4b* increase their expression during ripening with maxima reached at 55 DPA. Particularly, SlNADP-ME3 transcript abundance increased dramatically during pericarp development (400-fold higher in RR+7 compared to the EIG stage) (Figure 4).

**Figure 4.**
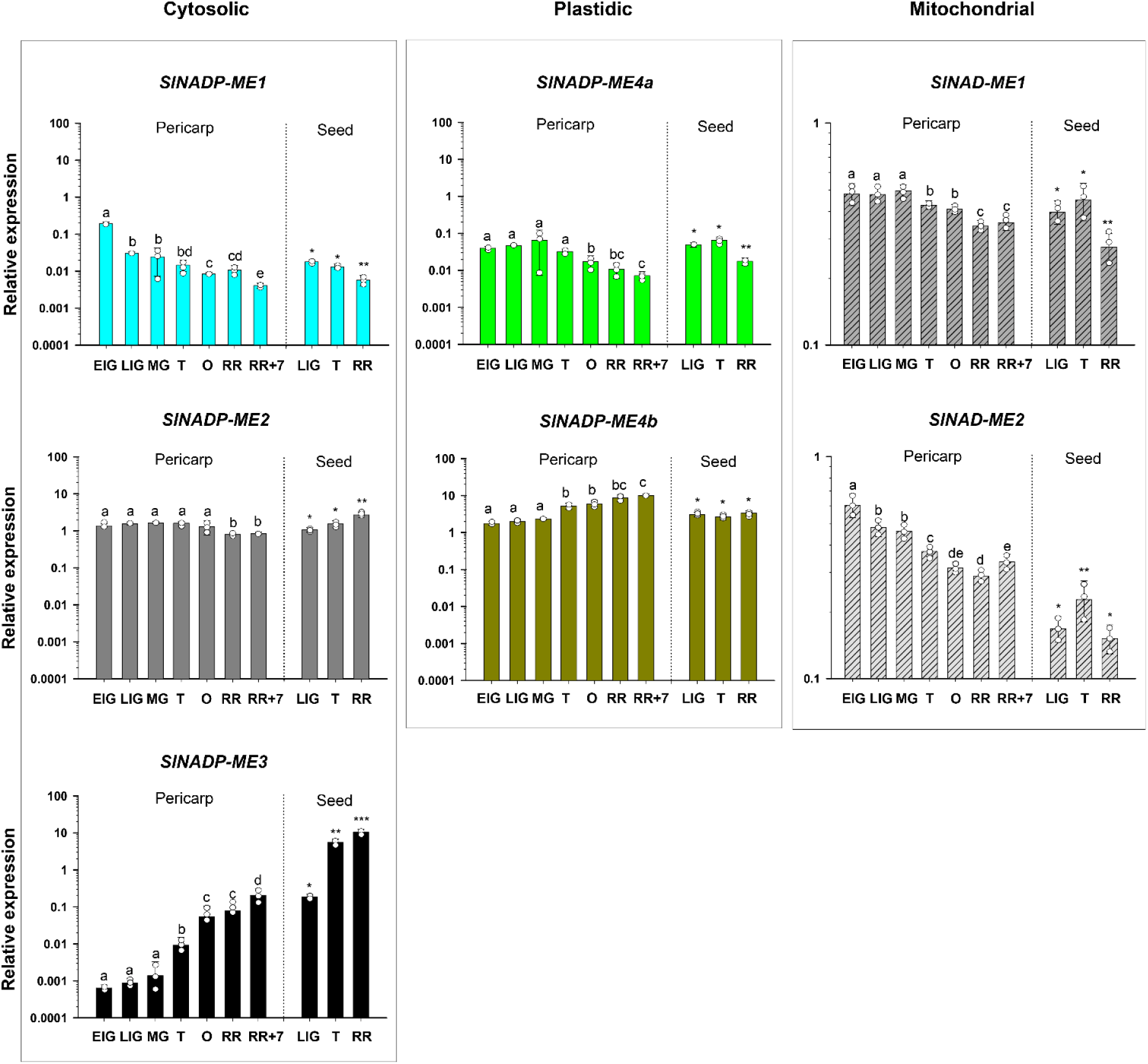
Expression levels of *SlNAD(P)-ME* genes throughout pericarp and seed development. Genes are presented in panels “Cytosolic”, “Plastidic” and “Mitochondrial” according to the subcellular location of the encoded protein. Transcript abundance was determined by qPCR and the values expressed relative to the abundance of the UBQ3 transcript. The y-axis is shown on a logarithmic scale. Fruits were harvested between 10 - 55 DPA (EIG - RR+7 stages) for **analysis**. Bars represent the mean ± s.d of three independent biological replicates (n=3). The same letters or dot (*) number on the bars indicate no statistical differences among stages based on one-way analysis of variance (ANOVA) for gene with Holm-Sidak method (p<0.05, n=3). The y-axis is shown on a logarithmic scale.

We also analyzed the *SlNAD(P)-ME* expression in seeds at three stage of fruit development: LIG (30 DPA), T (42 DPA) and RR (48 DPA), during which seed vigor (i.e., desiccation tolerance, germination percentage and longevity) is progressively increased (Bizouerne et al., 2021). We found that *SlNADP-ME3* is the most highly expressed gene in seeds (up to 10-fold higher than *UBQ3*), and its expression increases by over 60 times from the LIG to the RR stage (Figure 4). *SlNADP-ME2* is also largely up-regulated in seeds, where the accumulation of its transcript is ∼100-fold higher than that of *UBQ3* at the RR stage (Figure 4).

### Effect of conditions inhibiting ethylene-mediated responses on the expression profile of *SlNAD(P)-ME* genes

To explore the potential relationship between the expression profiles of *NAD(P)-ME* gene in fruit pericarp and ethylene, we treated fruits harvested at two stages of development: MG (38 DPA) and T (42 DPA) with 1-methylcylopropene (1-MCP), a known inhibitor of ethylene-dependent ripening responses. A delay in the progression of the color change was observed in 1-MCP-treated fruits (Figure S4A). Untreated MG and T tomatoes turned completely red after 6- and 10-days post-harvest (RR stage), respectively. However, treated T fruits remained in the O stage 6 days after treatment and treated MG fruits were between the O and RR stages after 10 days (Figure S4A). The expression level of *ACO6* (Solyc02g036350), which is ethylene-dependent and downregulated by 1-MCP (Bobokalonov et al., 2018), was significantly reduced in MG and T fruits incubated with 1-MCP as compared to untreated fruit (Figure S4B).

We analyzed Sl*NAD(P)-ME* genes in pericarp of 1-MCP-treated, untreated and control (immediately processed after harvest) fruits by qPCR (Figure 5). For cytosolic isoforms, we found that the expression level of *SlNADP-ME1* and *-2* genes was not significantly different between 1-MCP-treated and untreated samples (Figure 5A). However, *SlNADP-ME3* expression was enhanced by 1-MCP in MG-stage fruits after 10 days of treatment and in T-stage fruits (Figure 5A). Intriguingly, the NADP-ME3 transcript was not detected in the untreated MG fruits 4 days after harvest but accumulated at high levels in fruits that were incubated with 1-MCP.

**Figure 5.**
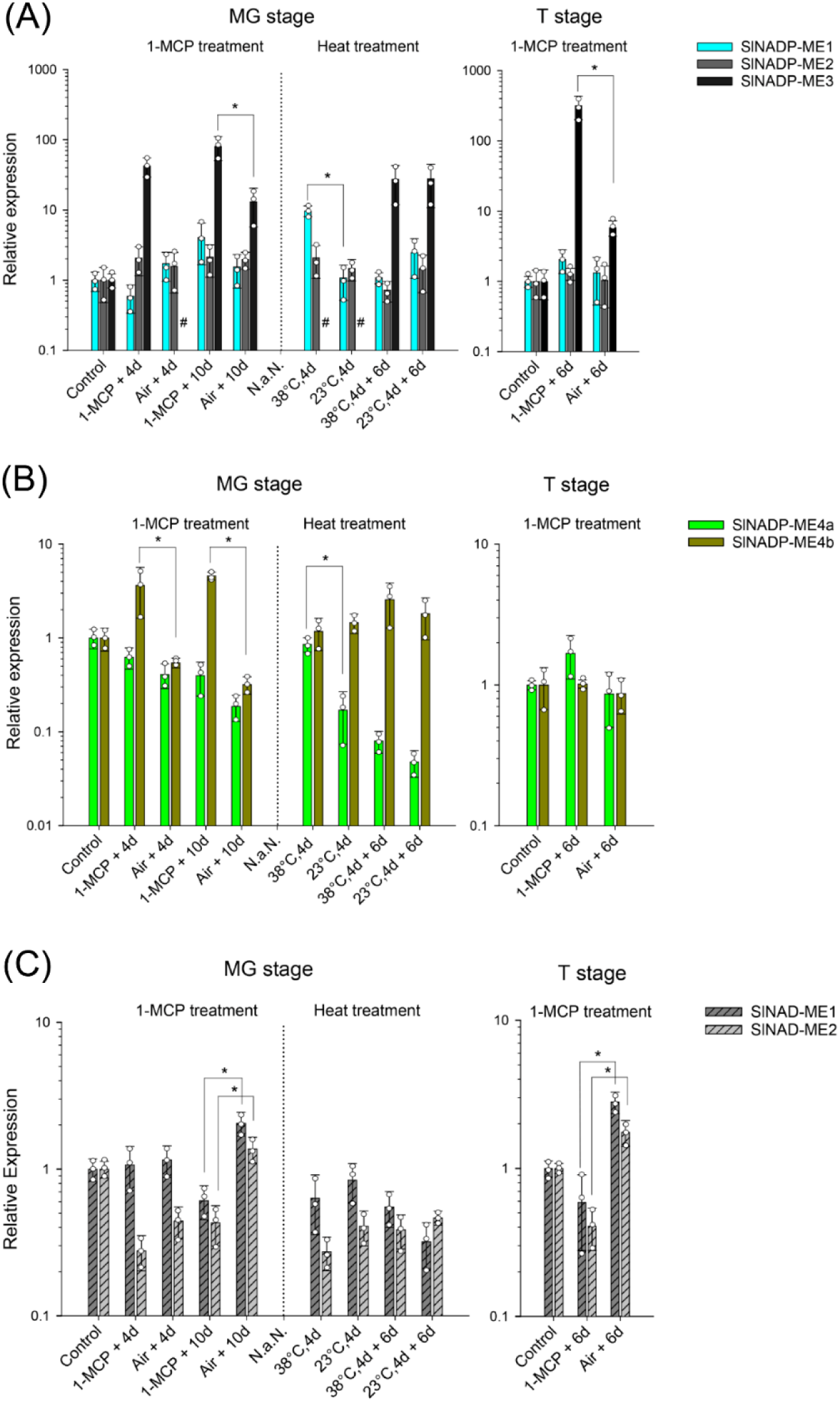
Expression levels of *SlNAD(P)-ME* in pericarp of fruits 1-MCP-treated or held at 38°C. Genes are grouped in different bar charts according to the subcellular location of the encoded protein: Cytosolic (**A**), Plastidic (**B**) and Mitochondrial (**C**). The levels of transcripts were assessed by qPCR and values expressed relative to the abundance in control (sample processed immediately after harvest). The y-axis is shown on a logarithmic scale. Bars represent the mean ± s.d of three independent biological replicates (n=3). Statistical differences (student’s t test, p<0.05, n=3) between treated and untreated samples are shown as *. Not detected transcript is indicated as #. Names defined for the samples on the x-axis: **1-MCP + 4d**, **6d** or **10d**: 1-MCP treated fruits (incubated with 1 ppm 1-MCP in closed chamber for 10 hours) and stored 4, 6 or 10 days, respectively, in the air in greenhouse before processing. **Air + 4d, 6d** or **10d**: untreated fruits (kept in closed chamber with air for 10 hours) and stored 4, 6 or 10 days, respectively, in the air in greenhouse before processing. **38°C** (or **23°C**)**,4d**: fruits held at 38°C (or 23°C) for 4 days before processing. **38°C** (or **23°C**)**,4d + 6d**: fruits held at 38°C (or 23°C) for 4 days and then transferred to 23°C in greenhouse for 6 days before processing (see Figure S4A).

Genes encoding for the plastid proteins, *SlNADP-ME4a* and *-4b*, were differentially affected by 1-MCP in MG-stage fruits (Figure 5B). In the presence of the inhibitor, the expression level of *SlNADP-ME4b* was about 4-fold higher than that of the untreated MG fruits at both sampled times, but not in fruits at the T stage (Figure 5B). No significant differences were observed in the level of expression of *SlNADP-ME4a* either in MG or T fruits by 1-MPC (Figure 5B). For mitochondrial heteromeric isoform, the expression of *SlNAD-ME1* and *-2* was inhibited by 1-MPC in MG fruits after 10 days of treatment and in T-stage fruits (Figure 5C).

Short exposures of fruits to temperatures above 35°C reversibly inhibit ethylene synthesis and tomato fruit ripening by downregulating the level of expression of *ACO* genes (Villalobos et al., 2011). Hence, we further analyzed the relation between expression of *NAD(P)-ME* genes and ethylene in tomato pericarp after heating. Fruits were harvested at the MG stage, placed at 38°C for 4 days and removed to 23°C for 6 days before analysis (Figure S4A). We found that the expression of the gene encoding the cytosolic SlNADP-ME1 and plastidic NADP-ME4a were about 10-fold higher in pericarp of MG fruits held at 38°C than in unheated fruits (Figure 5A). Nevertheless, when heated fruits were placed at 23°C for 6 days, the expression level of *SlNADP-ME1* and *-4a* were like unheated fruits. Intriguingly, at both temperatures tested, SlNADP-ME3 transcripts were not detected in MG fruits at 4 days after harvest but accumulated at high levels 10 days after harvest (Figure 5 A). The expression of the other *SlNAD(P)-ME* was not modified by temperature (Figure 5A and B). Finally, *ACO6* expression levels in heated fruit were 40 times lower than in untreated fruit (Figure S4B) and reached similar levels to the control when fruits were placed at 23° for 6 days, indicating a recovery of the heat-induced inhibition of ethylene biosynthesis.

### Identification of cis-acting regulatory elements in *SlNAD(P)-ME* promoters

To identify cis-acting regulatory elements (CAREs) in the promoter regions of each *SlNAD(P)-ME* gene, we conducted a sequence scan extending 1.5 kbp upstream and 100 pb downstream from the transcription start site using the PlantCare Supplementary Database (http://bioinformatics.psb.ugent.be/webtools/plantcare/html/). Our findings revealed a minimum of 29 CAREs associated with *SlNADP-ME4b* and *NAD-ME-2*, while *SlNADP-ME3* exhibited a higher number, with up to 42 CAREs identified (Supplementary Data 2). The cis-elements were categorized into five functional groups: (i) light responsive elements, (ii) environmental stress-related elements, (iii) hormone responsive elements, (iv) development related elements, and (v) site-binding related elements (Figure 6A and Supplementary Data 2).

**Figure 6.**
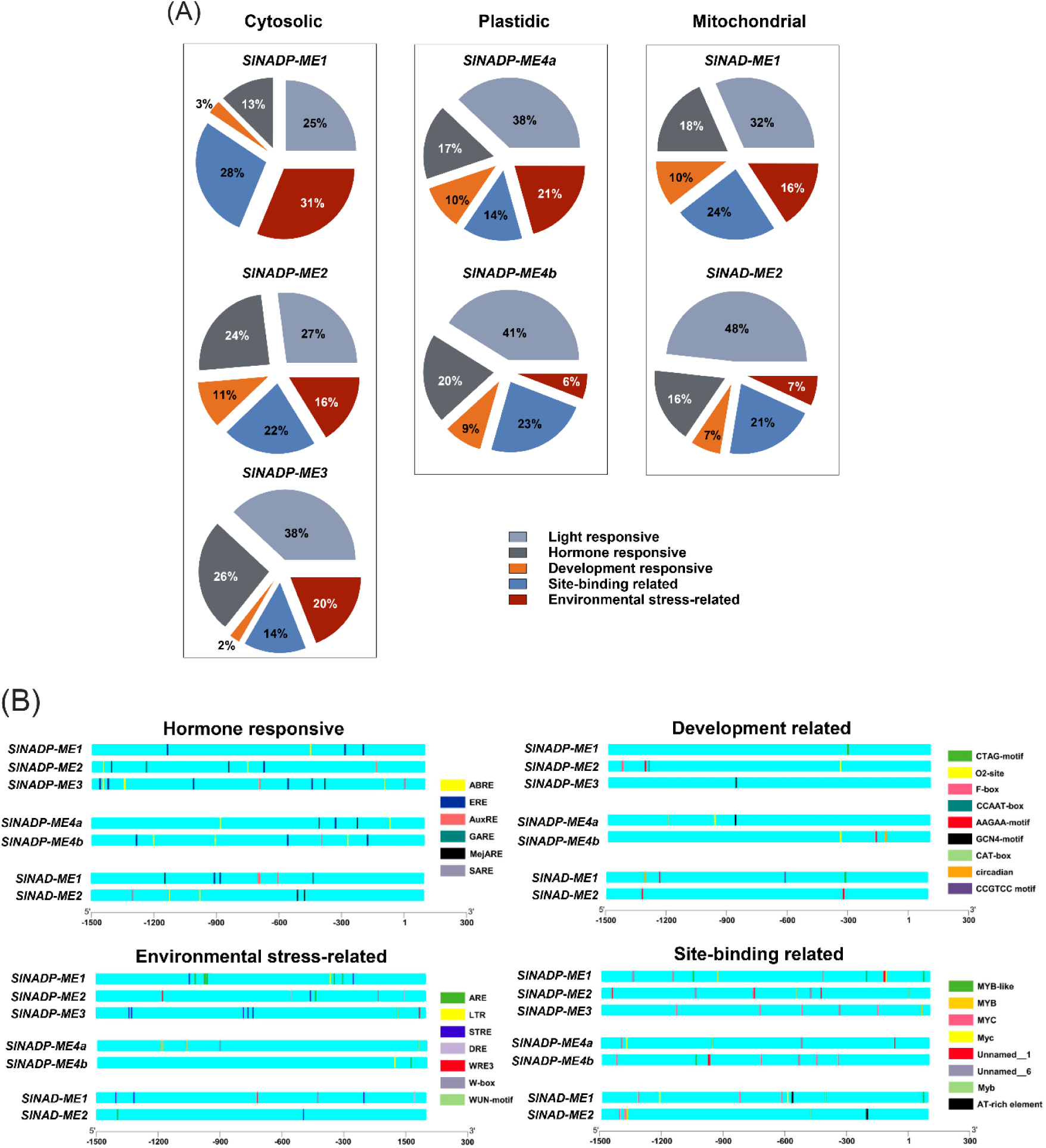
Cis-regulatory element analysis of *SlNAD(P)-ME* genes. (**A**) Pie distribution of identified motifs in *SlNAD(P)-ME* promoters based on their biological functions. For the calculation of the percentages, the number of times each element is present in each category per gene was considered (Date 2). (**B**) Position of identified cis-elements in *SlNAD(P)-ME* promoters. TSS (star transcription site) is numbered as 1.

*SlNADP-ME1* promoters show the highest proportion of environmental stress and binding site related elements compared to *SlNADP-ME2* and *-3* promoters (Figure 6A). In *SlNADP-ME3* light and hormone response categories together account for more than 60% of the total of regulatory sequences. In contrast, no significant bias in the proportion of the five categories of cis-elements in SlNADP-*ME2* was observed (Figure 5A).

Response elements to different environmental stresses and hormones are present in *SlNADP-ME1, -2 and -3* promoters (Figure 6B and Supplementary Data 2). *SlNADP-ME1* contains 6 ARE (cis-element for the anaerobic induction) copies, *SlNADP-ME2* shows 4 MejARE (methyl jasmonate acid responsive element), and *SlNADP-ME3* presents 5 copies of STRE (stress-responsive element), 5 ERE (ethylene responsive element), and 3 ABRE (abscisic acid responsive element).

The *SlNADP-ME2* promoter presents single copies of a variety of cis-elements related to development (endosperm-specific and meristem expression, and cell cycle regulation) and environmental stress (drought-inducibility and sites acting in biotic and abiotic stresses, seed dormancy, senescence, wounding and pathogen responses) (Figure 6B, Supplementary Data 2). These regulatory sequences are absent in cytosolic SlNADP-ME1 and -3 encoding genes. The *SlNADP-ME3* promoter contains a single developmental element: the GCN4-motif, a regulatory sequence involved in endosperm expression, and a diversity of light responsive elements such not found in other *SlNAD(P)-ME* promoters (Suppl. Figure 5 and Supplementary Data 2). In addition, *SlNADP-ME3* possesses cis-element for the major plant hormones: abscisic acid (ABA), ethylene, auxin, methyl jasmonate, and salicylic acid (SA).

*SlNADP-ME4a* and -4b promoters mostly differ in the cis-element related to environmental stress (21 % vs 6%) (Figure 6B, Supplementary Data 2). *SlNADP-ME4a* promoter possesses several copies of wounding responsive elements, and for maximal elicitor-mediated activation. On the other hand, *SlNADP-ME4b* presents one LRT motif (cis-acting element involved in temperature responsiveness) and one ARE (for anaerobic induction) copy.

Regarding genes encoding for the mitochondrial SlNAD-ME heteromer, *SlNAD-ME1* shows a higher number and diversity of cis-elements related to hormone (32% vs 16%) and environmental (19% vs 6%) responses than *SlNAD-ME2* (Figure 6A and B). This discrepancy in the promoter constitution of genes encoding a heteromeric enzyme could be overcome by considering that both promoters possess the AT-rich element (Figure 5B, Supplementary Data 2), a matrix-binding region that binds ATBP-1 (AT-rich DNA binding protein) and stimulates the recruitment of the molecular machinery of transcription (Tjaden et al., 1994). Thus, by tethering genes to nuclear scaffold, ATBP-1 could cover the spatial proximity of the *SlNAD-ME1* and *-2* promoters for the coordinate transcription of both *NAD-ME* genes.

### Biochemical characterization of recombinant SlNADP-MEs proteins

To gain deeper insights into the functional roles of the SlNADP-ME family, we conducted a comprehensive biochemical analysis of the recombinant forms of cytosolic SlNADP-ME1, -2, and -3, along with the mature plastidic proteins SlNADP-ME4a and -4b (lacking the targeting peptide). All isoforms exhibited enzymatic activity, although they displayed distinct biochemical characteristics. In native-PAGE assayed for NADP-ME activity, SlNADP-ME1 and -4b revealed similar protein band profiles, characterized by two major activity bands with differing mobilities (Figure 7A): a high mobility band that corresponds to an oligomeric state of a homodimer of approximately 146 kDa, and a low-mobility main band, suggesting the presence of larger assemblies, likely tetramers. Conversely, cytosolic SlNADP-ME2, -3 and plastidic SlNADP-ME4a showed a single activity band with low migration similar to the slower migration band of SlNADP-ME1 and -4b (Figure 7A and B), indicating the formation of large assemblies *in vitro.* It is important to highlight that, unlike the other isoforms, the activity band of SlNADP-ME3 in native PAGE exhibited very low intensity. This band only became visible after extended incubation times in the activity assay buffer, suggesting a unique behavior in terms of kinetic properties (Figure 7B). The differences in mobility observed between the plant tissue-derived and the recombinant protein (Figure 2 and 7) suggest that, in plant tissues, SlNADP-MEs may adopt a more native conformation, interact with other proteins in the extract, or undergo post-translational modifications beyond the capabilities of the prokaryotic expression system.

**Figure 7.**
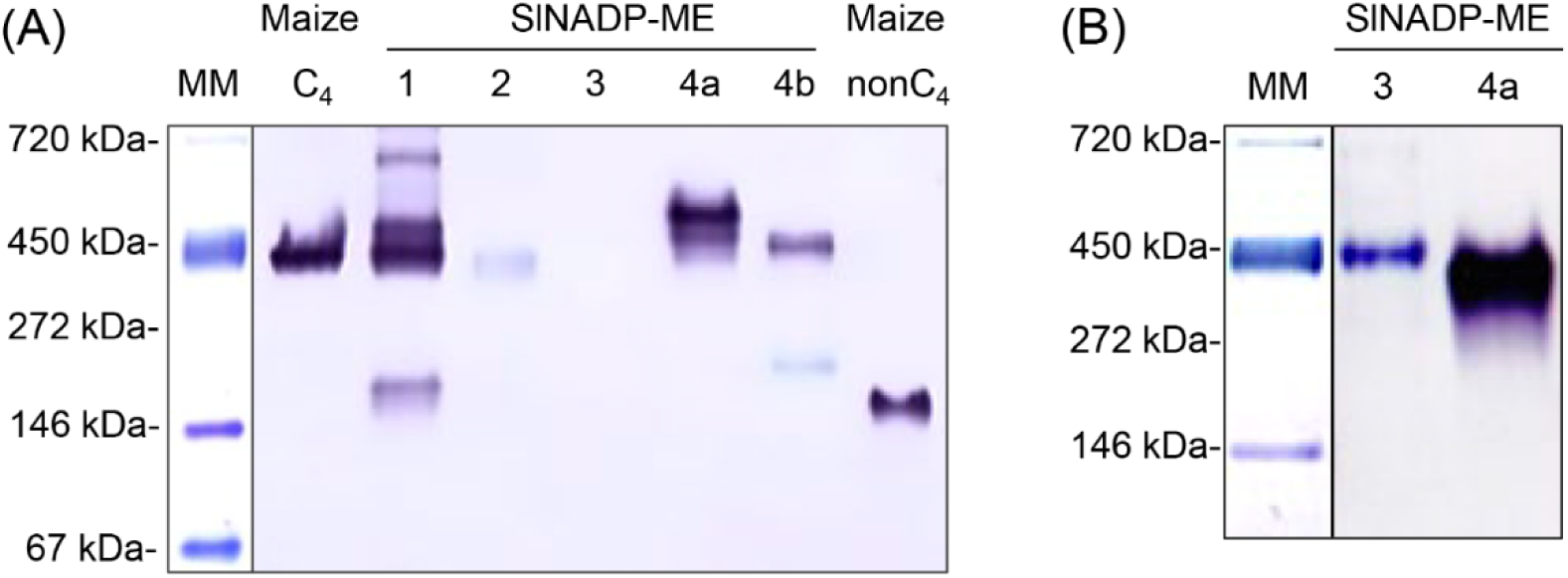
Native-PAGE of recombinant SlNADP-ME isoforms. Approximately 4 μg of each recombinant purified isoform was loaded. Lane 1, SlNADP-ME1; lane 2, SlNADP-ME2; lane 3, SlNADP-ME3; lane 4a, SlNADP-ME4a; lane 4b, SlNADP-ME4b. For the *in-gel* activity assays the gels were incubated for 15 min (**A**) or 60 min (**B**) in the assay medium at pH 7.5. A violet precipitate indicates NAD-ME activity. In (**A**) recombinant C4-NADP-ME and nonC4-NADP-ME from maize, which assemble as a tetramer and dimer, respectively, (Alvarez et al., 2019) were runs on the same gel. MM: Molecular mass marker.

For the forward reaction (malate oxidative decarboxylation), we found that SlNADPME4b exhibits the highest affinity toward NADP (K_m_∼10 μM), followed by SlNADP-ME1, -3 and -4a (K_m_∼20 μM), and SlNADP-ME2 with the lowest affinity (K_m_∼60 μM) (Table 1). Except for SlNADP-ME3, all isoforms have malate affinities (0.3 to 3 mM) similar to other plant NADP-MEs (Saigo et al., 2013; Alvarez at el., 2013; Garrard Wheeler et al., 2008). SlNADP-ME3 has the lowest malate affinity (K_m_∼10 mM). On the other hand, the cytosolic SlNADP-ME1 and -2 presented the highest specific activity values (k_cat_∼30 s^−1^ and ∼36 s^−1^, Table 1), followed by the plastid SlNADP-ME4a and -4b (k_cat_∼10 s^−1^ and ∼24 s^−1^, respectively). SlNADP-ME3 has a much lower specific activity, with a k_cat_ value 30-fold lower than that of the other cytosolic isoforms, SlNADP-ME1 and -2. SlNADP-ME4b has the highest catalytic efficiency (k_cat_/K_m_) for the plastidic isoforms (Table 1), while SlNADP-ME1 has the highest catalytic efficiency for the cytosolic isoforms and SlNADP-ME3 is a protein with an extremely low kinetic capacity *in vitro* (Table 1) in agreement with its behavior in native-PAGE (Figure 7).

**Table 1.** Kinetic properties of recombinant tomato NADP-ME isoforms.The indicated values, obtained by nonlinear regression, are the average of measurements ± SE with at least three different batches of recombinant enzymes.

|  | SINADP-ME1 | SINADP-ME2 | SINADP-ME3 | SINADP-ME4a | SINADP-ME4b |
| --- | --- | --- | --- | --- | --- |
| **Malate oxidative decarboxylation** |  |  |  |  |  |
| $K_m \text{ NADP } (\mu\text{M})$ | $17.7 \pm 1.8$ | **$56.7 \pm 3.6$** | $16.6 \pm 3.6$ | $22.3 \pm 3.7$ | $11.3 \pm 2.8$ |
| $K_m \text{ malate } (\text{mM})$ | $0.47 \pm 0.02$ | $2.35 \pm 0.24$ | **$9.60 \pm 3.83$** | $0.59 \pm 0.08$ | $0.42 \pm 0.03$ |
| $k_{\text{cat}} (\text{s}^{-1})$ | $30.1 \pm 0.7$ | $35.7 \pm 0.7$ | **$1.8 \pm 0.2$** | $11.2 \pm 0.3$ | $23.8 \pm 0.8$ |
| $k_{\text{cat}}/K_m \text{ NADP } (\text{s}^{-1} \mu\text{M}^{-1})$ | 1.70 | 0.63 | **0.11** | 0.50 | 2.11 |
| $k_{\text{cat}}/K_m \text{ malate } (\text{s}^{-1} \text{mM}^{-1})$ | 64.0 | 15.2 | **0.20** | 19.0 | 56.6 |
| **Pyruvate reductive carboxylation** |  |  |  |  |  |
| $K_m \text{ pyr } (\text{mM})$ | $22.1 \pm 0.7$ | $5.0 \pm 1.1$ | - | $15.5 \pm 1.0$ | $1.1 \pm 0.3$ |
| $k_{\text{cat}} (\text{s}^{-1})$ | $15.3 \pm 2.0$ | $26.5 \pm 1.7$ | - | $5.3 \pm 0.8$ | $22.3 \pm 1.4$ |
| $k_{\text{cat}}/K_m \text{ pyr } (\text{s}^{-1} \text{mM}^{-1})$ | 0.7 | 5.3 | - | 0.34 | 20.2 |
| **Relation forward/reverse reaction** |  |  |  |  |  |
| $k_{\text{cat}}/K_m \text{ malate/}$<br>$k_{\text{cat}}/K_m \text{ pyr}$ | 91.4 | 2.8 | - | 56.0 | 3.0 |

For the reverse reaction (pyruvate carboxylation), plastidic SlNADP-ME4b and cytosolic SlNADP-ME2 exhibited the highest catalytic efficiencies (k_cat_/K_m_ ∼20 **s^−1^mM^−1^** and ∼5 **s^−1^mM^−1^**, respectively) (Table 1) with specific activity values comparable to those for the direct reaction (k_cat_∼22.3 s^−1^ and ∼26.5 s^−1^) and high pyruvate affinity (K_m_ ∼1 mM and ∼5 mM) (Table 1). Thus, unlike other isoforms, which showed lower k_cat_ values and/or pyruvate affinities for the carboxylation reaction (Table 1), SlNADP-ME4b and SlNADP-ME2 are enzyme with comparable decarboxylase/carboxylase activity *in vitro* (Table 1).

## DISCUSSION

### Phenotypic similarity of expression of NAD(P)-MEs family members in major tissues of eudicot plants

In eudicot, the minimal constitution of the NAD(P)-ME family comprises two cytosolic isoforms, one plastid protein and two mitochondrial subunits (Tronconi et al., 2018). Most species exhibit variable expansion in NAD(P)-ME members as consequence of developmental, physiological and/or environmental particular requirements. *S. lycopersicum* encoding five SlNADP-ME proteins that are distinct in their tissue localization and biochemical properties. Cytosolic SlNADP-ME1 and -2 are present in all major tissues of flowering plants, as are the two plastid isoforms SlNADP-ME4a and -4b (Figure S2). Furthermore, SlNADP-ME2 and -4b would be housekeeping proteins involved in the ontogenetic processes of plant physiology, according to the substantial accumulation of their transcripts compared to those for SlNADP-ME1 and -4a (Figure S2).

The *SlNADP-ME2* promoter contains several developmentally related boxes (Figure 6, Supplementary Data 2) and the encoded enzyme has similar kinetic parameters (Table1) than AtNADP-ME2 (Gerrard Wheeler et al., 2005), Thus, this highly expressed protein could participate in the generation of reducing power, assisting the oxidative pentose phosphate pathway, under biosynthetic functions requiring high levels of NADPH in cytosol, as was proposed for AtNADP-ME2 homolog (Maurino et al., 2009; Gerrard Wheeler et al., 2005). In turn, cytosolic SlNADP-ME1 would be mainly present in defense and antioxidant mechanisms, given the large proportion of stress-related *cis*-elements in the promoter, and its expression specifically increased in fruits held at 38°C (Figure 5A and 6, Supplementary Data 2). In line with this, *SlNADP-ME1* is up-regulated, to provide NADPH for ROS synthesis, during systemic resistance responses triggered by *Oidium neolycopersici* infection (Pei et al., 2011), and in leaves infected with *Pseudomonas syringae* (http://ted.bti.cornell.edu ). Therefore, it appears that the proposed biosynthetic and defense functions of cytosolic AtNADP-ME2 from *A. thaliana* (Voll et al., 2012) are conducted by two constitutively co-localized SlNADP-ME1 and -2 isoforms with distinct kinetic properties in *S. lycopersicum*. Conversely, cytosolic *SlNADP-ME3* expression is highly constrained in tissues, limited almost exclusively to the roots of flowering plants. *SlNADP-ME3* promoter contains the highest proportion of *cis*-elements for plant hormones, mainly ABA and ethylene (Figure 6, Supplementary Data 2), indicating tight gene regulation through a complex crosstalk of internal and external signals intended for their participation in specialized processes. This pattern of gene expression and regulation resembles that described for AtNADP-ME1(Arias et al., 2018; Yazdanpanah et al., 2018; Badia et al., 2020), thus, SlNADP-ME3 could also be important in ABA-mediated water deficit responses and Al^+3^ stress tolerance in roots.

On the other hand, the presence of two plastidic NADP-ME isoforms is a common property of some C_4_-plants (Tausta et al., 2002; Saigo et al., 2004; Christin et al., 2009; Alvarez et al., 2019). However, we found that *S. lycopersicum* possesses two functional plastidic-encoding *SlNADP-ME* genes. *SlNADP-ME4a* is expressed 100-fold less than *SlNADP-ME4b* (Figure S2), but its expression specifically increased in fruits held at 38 °C (Figure 5B and 6) and the *cis*-element composition of its promoter suggests that SlNADP-ME4a may be specialized in defense processes against environmental stress (Figure 6 and Supplementary Data 2). On the other hand, SlNADP-ME4b could play a key role in fat biosynthesis, as evidenced by its expression pattern and substrate affinity that closely resemble those of the plastidic AtNADP-ME4 (Table 1, Gerrard Wheller et al., 2005), which has been proposed to be involved in malate-dependent lipid biosynthesis in fast-growing tissues (Preiss et al., 1994; Maurino et al., 2009).

Therefore, a tighter functional specification of SlNADP-ME isoforms emerges as distinctive property of tomato due to processes of gene specialization (SlNADP-ME1) and sub-functionalization (SlNADP-ME4a and -4b). Regardless of the phylogenetic relationship between SlNADP-ME and AtNADP-ME members, the significant similarity in biochemical properties and expression pattern observed in the major organs of flowering plants suggests that this family of enzymes plays a role in the morphological program of these two distantly related species, most likely because of adaptive selection that shaped these phenotypic similarities.

### Distinctive expression pattern of *NAD(P)-ME* genes during tomato fruit development suggests specific roles for encoded proteins

NADP-ME activity increased slowly over the course of the early stages of pericarp development (Figure 1D) but then abruptly increased and reached a plateau at the MG stage. The appearance of a NADP-ME isoform of high mobility observed in native-PAGE (Figure 2B) during the cell expansion phase could be associated with the initial activity profile (Figure 1D).

Cell expansion and ripening phases in tomato pericarp are well defined by the large changes in the activity of a substantial set of enzymes involved in primary and secondary metabolism (Steinhauser et al., 2010; Biais et al., 2014). Plant NADP-ME may play anabolic or catabolic roles depending on the physiological context and the isoform involved (Badia et al., 2017). In anaplerosis (pyruvate carboxylation reaction), SlNADP-ME would function in coordination with PEPC by restoring the reserves of malate and OAA used for amino acid biosynthesis and providing the osmotic driving force governing cell expansion (Carrari and Fernie, 2006; Beauvoit et al., 2014). This hypothesis is supported by the significant carboxylase activity shown by SlNADP-ME2 and -4b (Table 1). On the other hand, as a decarboxylating enzyme, SlNADP-ME might be providing reducing power for starch and lipid synthesis in plastid. The ADP-Glc pyrophosphorylase (AGPase) participates in starch synthesis and is activated by the reduction of cystine residues (Centeno et al., 2011). AGPase activity is highest at the beginning of tomato cell expansion and remains high until the MG stage (Osorio et al., 2013). Thus, in a coordinated activity profile of these two enzymes in cell expansion phases, the redox activation state of AGPase could be modulated by the amount of NADPH dispensed by a SlNADP-ME isoform. Consistent with this, transgenic tomato plants with reduced levels of plastid NADP-ME activity had lower starch levels in fruit at the MG stage due to decreased redox activation of AGPase (Centeno et al., 2011; Osorio et al., 2013). Additionally, lipid synthesis for new membrane deposition in fruit cell growth demands NADPH and pyruvate, which would also be provided by the plastidic SlNADP-ME (Smith et al., 1992; Kang et al., 1994; Preiss et al., 1994).

As mentioned before, SlNADP-ME4b could support malate-dependent lipid biosynthesis in fast-growing tissues. Several regulatory sequences for auxin and ABA, the key hormones governing fruit cell division and expansion, are present in the *SlNADP-ME4b* promoter (Supplementary Data 2), as well as multiple ethylene-dependent cis-regulatory elements. Hence, at the first stage of cell expansion, auxins delivered by the seeds moderately increase SlNADP-ME4b activity (Obroucheva et al., 2014). Then, ABA levels increase in fruit (Zhang et al., 2009; Zhang et al., 2023) and strongly enhances the levels of SlNADP-ME4b activity, which provide NADPH and pyruvate required at the last phase of cell expansion. In the terminal stage of fruit growth, ethylene acts as an inhibitor of *SlNADPME4b* expression (MG stage, Figure 5), restricting ABA-dependent cell growth effects. With ABA signaling declining, the forward transition of fruit ripening occurs, and the dependence of *SlNADP-ME4b* on ethylene ends (T stage, Figure 5).

During fruit ripening phase, we did not find any straightforward relationship between changes in NADP-ME activity and transcripts of the encoded enzymes (Figure 1D and Figure 4). This decoupling between transcript accumulation and protein synthesis/degradation may be due to the different phases of fruit development being highly dynamic and involving processes such as cell differentiation/ specialization or stress responses, where posttranscriptional processes may lead to strong deviations from an ideal correlation (Liu et al., 2016; Bias et al., 2014; Steinhauser et al., 2010). Hence, high *NAD(P)-ME* expression could still occur in specialized pericarp cells during maturation. According to the Tomato Expression Supplementary Data (TEA, https://tea.solgenomics.net), cytosolic SlNADP-ME2 and plastid SlNADP-ME4b are expressed in the outer epidermis, exocarp, and inner epidermis during all phases of tomato development. These tissues have a highly active secondary metabolism responsible for outer and inner cuticle synthesis, carotenoids, flavonoids, phenolics, and volatile compounds. Malate-mediated ripening control of changes in pigmentation or postharvest characteristics is conducted primarily by mitochondrial enzymes (e.g., fumarase) and then, to a lesser extension, by cytosolic and/or plastidic SlNADP-MEs (Centeno et al., 2011; Osorio et al., 2013).

SlNADP-ME2 and -4b are also expressed in mature tomato seeds. In Arabidopsis, cytosolic AtNADPME2 and plastidic AtNADP-ME4 homologs are highly present in the endosperm (Gerrard Wheeler et al. 2005). Thus, the high level of expression of SlNADP-ME2 in tomato seeds could be correlated with the generation of reducing power for anabolic processes, assisting the oxidative pentose phosphate pathway in the cytosol. In addition, plastidic SlNADP-ME4b probably not only participates in the demand for new membrane synthesis during fruit growth, but also in lipid deposition in the endosperm (Preiss et al., 1994; Smith et al., 1992).

In fruit, *SlNADP-ME3* expression is mainly restricted to seeds where its transcript accumulates at high levels with ripening (Figure 4), which is consistent with the presence of the GCN4 motif in its promoter region (Figure 6, Supplementary Data 2). In Arabidopsis, accumulation of the AtNADP-ME1 ortholog (Figure 3; Gerrard Wheeler et al. 2005) in seeds is critical to protect against oxidation during dry storage, preserving their longevity and dormancy (Yazdanpanah et al., 2018). In fact, carbonylation of seed proteins is enhanced in *nadp-me1 knockout* lines, indicating that AtNADP-ME1 acts avoiding ROS excessive levels. For successful germination, an optimal level of ROS is required (Bailly et al., 2008). Several carbonylated forms of AtNADP-ME1 were identified in seeds (Galland et al., 2012). A highly carbonylated AtNADP-ME1 was present in seeds that lost their germination capacity (Rajjou et al. 2008). Thus, high carbonylation would result in reduced AtNADP-ME1 activity and excess ROS. However, lower carbonylation states of AtNADP-ME1 could stimulate enzymatic activity, providing NADPH to reach optimal ROS levels. Using the iCarSP software (Zhang et al., 2021; http://lin-group.cn/server/iCarPS) we found that SlNADP-ME3 and AtNADP-ME1, but not the other NADP-ME members, are carbonylated in 167K residue. Alphafold 3 modeling of SlNADP-ME3 (Figure S6) shows that the 167K-semialdehyde (K-carbonyl form) modifies the conformation of the helical segment that contains the 170D residue, only present in AtNADP-ME1 and SlNADP-ME3 (Figure S6), which establishes an ionic bond with 589R in the other dimer of the tetramer. Recently, it has been shown that interactions between C-terminal residues at the dimer interface are crucial for proper oligomeric stability and enzymatic functionality in plastidic NADP-MEs from Poaceae (Böhm et al., 2025). Therefore, changes in dimer-dimer contacts by K-carbonylation in SlNADP-ME3 could be an ROS-sensing regulatory mechanism that improves its catalytic efficiency (Table 1) for normal seed germination. In line with this, K-carbonylation has been shown to be essential to activate the peroxidase activity of cytochrome *c* during cell apoptosis (Yin et al. 2019), and histone H1 K-carbonylation influence the compaction of chromatin and the recruitment of trans-acting factors (Izzo and Schneider, 1016).

ROS homeostasis function of SlNADP-ME3 might be also required in the pericarp of mature red fruits (RR+7), where it accumulates at the highest levels (Figure 4). Mitochondrial SlNAD-ME activity remains high (75%) during the later stages of ripening (Figure 1), where heat-coupled malate respiration could take place to ensure the development of desirable fruit properties, and ROS levels could become a harmful by-product. Our results indicate that ethylene maintains malate-related mitochondrial respiration during ripening, as genes encoding heteromeric SlNAD-ME were inhibited during 1-MCP treatment (Figure 5C).

### The common program of NAD(P)-ME families in eudicot was traced through multiple evolutionary trajectories

It may be a common trait for eudicot plant species to possess two *NADP-ME* genes constitutively expressed in photosynthetic and non-photosynthetic cells, one providing high enzymatic activity in the cytosol (e.g. SlNADP-ME2 and AtNADP-ME2) and the other in the plastids (e.g. SlNADP-ME4b and AtNADP-ME4). The functional similarity of NADP-ME in cytosol and plastid from distant species is supported by the very similar kinetic properties shown by recombinant isoforms of *S. lycopersicum* (Table 1) and *A. thaliana* (Gerrad Wheeler et al., 2005, 2008). Despite this, the molecular mechanisms that led to this phenotypic similarity in protein expression, biochemical properties and function are not necessarily due to phylogenetic gene relationships (Cerca, 2023). In this sense, *SlNADP-ME2* and *AtNADP-ME2* belong to different phylogenetic clades with very high confidence (Figure 3 and Figure S1). Furthermore, *SlNADP-ME2* presents an independent genomic configuration that is specific to Solanaceae (Figure S2), probably originating from Solanaceae-specific evolutionary event. After gene duplication originating *SlNADP-ME2* and *-3* paralogs, chromosomal rearrangements in a Solanaceae ancestor (Figure S2) likely allowed *SlNADP-ME2* to evolve by modifying its tissue expression and biochemical properties from those defined from its ancient phylogenetic origin (Lineage I; Figure 3 and Figure S1). That said, convergent evolution is the more parsimony explanation for phenotypic similarity between SlNADP-ME2 and AtNADP-ME2 (Figure 8).

**Figure 8.**
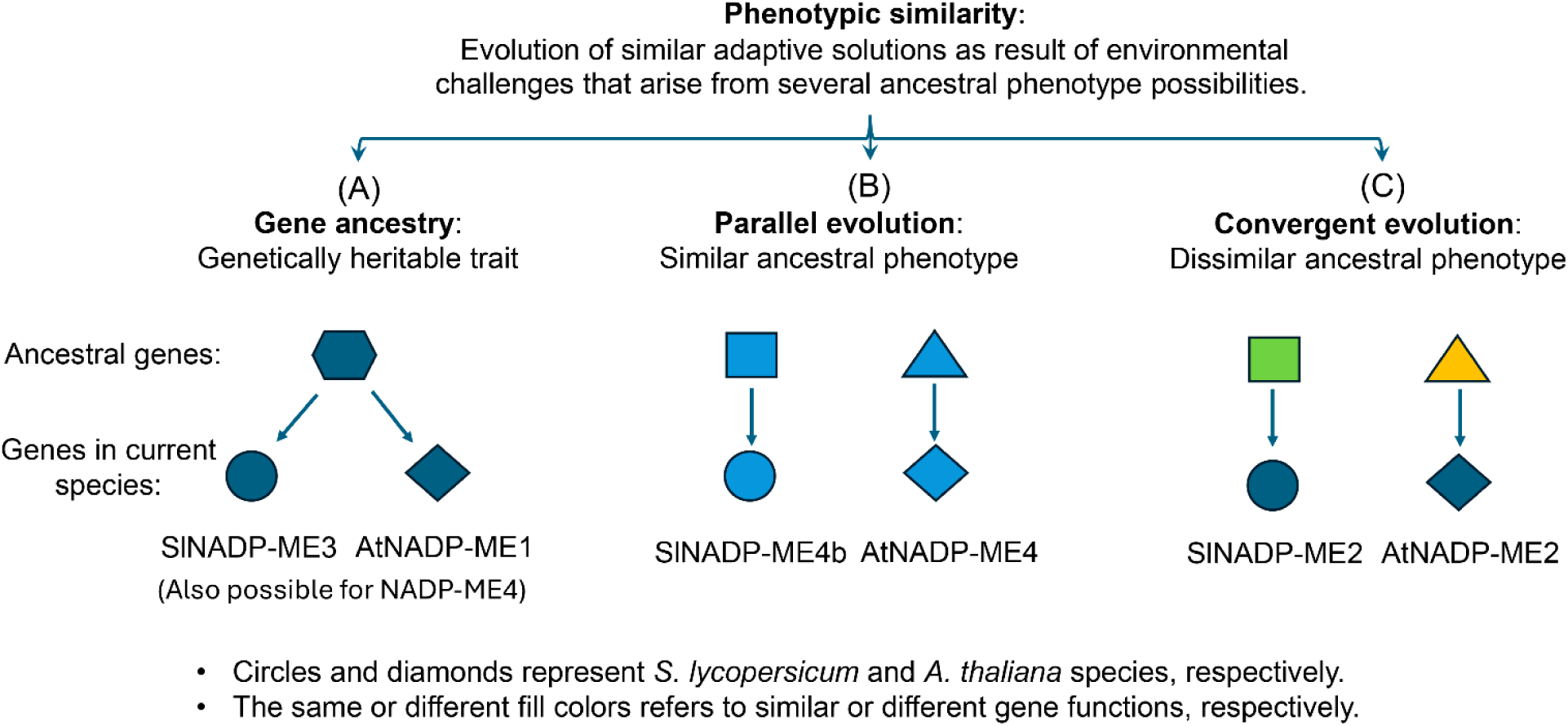
(**A**) Genetic ancestry refers to the lineage or heritage of an individual, tracing back the genetic line through generations to understand how certain genetic traits or markers have been passed down. (**B**) Parallel gene evolution refers to the phenomenon where similar traits or genes evolve independently in different species due to similar environmental pressures, rather than from a common ancestor. (**C**) Genetic convergence refers to evolution of a similar adaptive solution in response to environmental challenges from distinct ancestral phenotypes. Figure adapted from Cerca (2023).

On the other hand, the widespread tissue expression of the genes encoding plastidic *NADP-ME4b* may be a trait inherited from an ancient common ancestor with AtNADP-ME4 (Figure 3 and Figure S1) although it cannot be ruled out that it is the result of a parallel evolution phenomenon, which is common in distant species (Figure 8; Cerca, 2023). Interestingly, SlNADP-ME3 and AtNADP-ME1 proteins: (i) share a distinct expression pattern in relation to the other NADP-MEs, (ii) are the isoforms with the lowest enzymatic efficiency in both plant species, and (iii) would be specifically regulated by carbonylation. With this in mind, we suggest that these unique properties exhibited by SlNADP-ME3 and AtNADP-ME1 were delineated in an ancient common ancestor that possibly even predates the origins of eudicots (Figure 3, Figure S1; Gerrard Wheeler et al., 2005; Alvarez et al., 2013; Arias et al., 2018). Thus, such foundational functions, possibly common to all angiosperms, were not acquired by repeated evolutionary events, but respond to a long-conserved and inherited genetic lineage (Figure 8).

To conclude, the significant phenotypic similarity in the biochemical properties and expression pattern of NAD(P)-ME members in two distantly related plant species, such as tomato and Arabidopsis, appears to have been markedly shaped by repeated evolution acting on different genetic sources (Figure 8).

## MATERIALS AND METHODS

### Pant material, growth conditions and fruit treatments

Seeds of the *Solanum lycopersicum* Micro-Tom variety were germinated into 1-L individual pots. The posts were first placed at 30°C for 72 h and then transferred to a growth chamber with a light flux of 110 ± 10 µE m^−2^ s^−1^ under a 16h/8h light/dark regime and at 23 ± 3 °C temperature. After flowering, the plants were trimmed to allow the development of four fruits bunches, on the second and thirst truss, per plant. Individual flowers were tagged at the anthesis phase, and the stages of fruit development were defined according to the days post anthesis (DPA). Upon harvesting, each fruit was cut into two parts, and the pericarp was separated from the placental tissue. Then, pericarp was cut into slices (of approx. 50 mg) and 3 to 5 slices were ground to a fine powder under liquid nitrogen and stored at -80° C until use. One ground powder constitutes a biological sample, and 3 to 5 biological samples (*n* replicates) were used for analyses.

For heat treatment, mature green fruits were obtained directly from a greenhouse and divided into three batches (Figure S4A). The first batch was immediately processed and stored (control). The second was placed at 23°C for 4 and 10 days. The third batch was heated in a chamber at 38°C for 4 days; then, half of the batch was removed, and the rest was placed for 7 more days at 23°C (Figure S4A). Relative humidity was maintained at 90 to 95%. At each time assayed, fruit pericarp from at least 5 fruits was processed as described above.

For the 1-MCP treatment, mature green and turning fruit were obtained from the same lot of plants as for the heat treatment and divided into three batches (Figure S4A). The first batch was immediately processed and stored (control). For mature green fruits, the second batch was treated with the inhibitor of ethylene perception (cc = 1 ppm) in a hermetic container for 10 h; then, half of the batch was stored for 4 days at 23°C, and the rest was placed for 6 more days at 23°C. The third batch was kept in the same conditions without 1-MCP; then, half of the batch was stored for 4 days at 23°C, and the rest was stored for 6 more days at 23°C. For turning fruits, the second batch consisted of fruits treated with 1-MCP (cc = 1 ppm) in a hermetic container for 10 h and then stored for 6 days at 23°C. The third batch was kept in the same conditions without 1-MCP and then stored for 6 days at 23°C (Figure S4B). At each time assayed, fruit pericarp from at least 5 fruits was processed as described above.

### Sample extractions

For enzymatic assays and non-denaturing PAGEs, the powders were treated by vigorous shaking with a 2/1 (ml/mg) ratio of ice-cold extraction buffer composed of 50 mM MES pH 6.5, 10 mM MnCl_2_, 1 mM EDTA, 10 mM 2-mercaptoethanol, 2% (v/v) glycerol, 0.5% (v/v) Triton X-100, 1 mM phenylmethylsulfonyl fluoride (PMSF) and polyvinylpyrrolidone (PVPP) in the presence of a protease inhibitor cocktail (Sigma). Homogenates were centrifuged for 20 min at 12 000 g and 4°C, and clarified extracts were desalted in Sephadex G-50 columns equilibrated with 50 mM Tris-HCl pH 8, 20% (v/v) glycerol and 5 mM 2-mercaptoethanol.

For denaturing PAGEs, 2 g of powder tissue were treated with 8 ml of extraction buffer composed of 0.1 M Tris-HCl pH 8.8, 10 mM EDTA, 0.9 M sucrose, 0.4% (v/v) 2-mercaptoethanol, 1 mM PMSF and 10 ml of phenol saturated with ice-cold TRIS-HCl, pH 8.8, and then agitated at 4°C for 30 min. The aqueous phases were back-extracted with extraction media and phenol by shaking. Tubes were centrifuged at 5000 g for 15 min at 4°C and the phenolic phases were transferred to a new tube leaving the interface intact. Proteins were precipitated with 5 vols of cold 0.1 M ammonium acetate in methanol at -20°C overnight. The samples were collected by centrifugation for 20 min at 14000 g and 4°C. Next, the pellet was washed with 1.5 ml of cold ammonium acetate/methanol and twice with cold 80% (v/v) acetone. One final wash used 1.5 ml of cold 70% (v/v) ethanol. Finally, the pellet was resuspended in 0.25 M Tris-HCl, pH 7.5, 2% (w/v) S.DS, 0.5% (v/v) 2-mercaptoethanol, and 0.1% (v/v) bromophenol blue, and boiled for 2 min.

Total leaf extracts and mitochondria leaf extracts were obtained as described in Tronconi et al. (2008).

### NAD(P)-ME activity assays

The activity of the enzymes was measured spectrophotometrically in a final volume of 1 ml at 30°C and 340 nm (ε_340nm_= 6.22 mM-1cm-1) using a UNICAM Helios b spectrophotometer (UNICAM Instruments, Cambridge, UK). The reaction mixtures used for each enzyme were as given below.

NADP-ME: 50 mM Tris-HCl, pH 7.5, 0.5 mM NADP; 10 mM malate, 10 mM MgCl_2_ and 10 to 30 µl of extract. The reaction started with malate (Detarsio et al., 2003).

NAD-ME: 50 mM MES-NaOH, pH 6.5, 2 mM NAD, 20 mM malate, 10 mM MnCl_2_ and 1 U of malate dehydrogenase (MDH). After the rapid increase in absorbance, 80 µl of extract were added and the subsequent steady increase was attributed to the decarboxylation of malate by NAD-ME (Tronconi et al., 2008).

Initial velocity studies were performed by varying the concentration of one of the substrates around its K_m_ value while keeping the other substrate concentrations at saturating levels at pH7.5. All kinetic parameters were calculated at least by independent triplicate determinations and adjusted to nonlinear regression using free concentrations of all substrates (Detarsio et al., 2003). One unit is defined as the amount of enzyme that catalyzes the formation of 1 mmol of NADPH min^−1^ under the specified conditions.

### Protein concentration measurements

Protein concentration was determined in crude extracts by the method of Bradford (1976) using the Bio-Rad protein assay reagent (Bio-Rad, Hercules, CA, USA) and bovine serum albumin as standard.

### Gel electrophoresis

SDS-PAGE was performed in 10% (w/v) or 7.5% to 10% (w/v) linear gradient polyacrylamide gels according to Laemmli (1970). Proteins were visualized with Coomassie Blue or electroblotted onto a nitrocellulose membrane for immunoblotting. Antibodies against maize recombinant nonC4-NADP-ME (Saigo et al., 2004), Arabidopsis recombinant NAD-ME1 and -2 (Tronconi et al., 2008), the major subunit of purified spinach RuBisCO large subunit (RuBisCO LSU, provided by Dr A Viale; National University of Rosario, Argentina) and the purified PEPCK from pineapple (Martin et al., 2011) were used for detection (1:100 to 1:400). Bound antibodies were visualized by linking to alkaline phosphatase-conjugated goat anti-rabbit IgG according to the manufacturer’s instructions (Sigma). Alkaline phosphatase activity was detected colorimetrically or by using a chemiluminescent kit (Immun-Star; Bio-Rad).

Native-PAGE was performed using a 6% (w/v) acrylamide separating gel. Electrophoresis was run at 150 V at 10°C. Gels were assayed for NADP-ME activity by incubating the gel in a solution containing 50 mM Tris-HCl, pH 7.5, 30 mM malate, 10 mM MgCl_2_, 0.8 mM NADP, 35 mg/mL nitroblue tetrazolium, and 0.85 mg/mL phenazine methosulfate at 30°C.

### RNA extraction and reverse transcription reaction

Total RNA was isolated from 50-100 mg of fruit pericarp, leaves, stems, flowers, seeds and roots using the Trizol reagent (Invitrogen) following the recommendations of the manufacturer. The concentration and integrity of the preparations were assayed by agarose 2 % gel electrophoresis. Two micrograms of total RNA were first treated with 1 U of RQ1 DNase (Promega) and, then, reverse transcribed with 200 U of M-MLV reverse transcriptase (Promega) using oligodT as primer. The cDNAs were used as templates for quantitative real-time PCR (qPCR) assays.

### Quantitative real-time PCR analysis

Relative gene expression was determined by qPCR on a Mx3000P detection system and the MxPro–Mx3005P software version 4.01 (Stratagene). Primers were designed using Primer Select of DNAStar Lasergene version 10.1 (DNASTAR, Madison, WI, USA) to allow for amplification of 150–250 bp products of similar GC and Tm characteristics. The generation of specific PCR products was confirmed by melting curve and gel analysis. Each qPCR reaction (20 µl) contained SYBR^®^ Green I 0.5 X (Invitrogen); 3 mM MgCl2; 0.5 µM of each primer; 0.5 U Platinum Taq, 1 X buffer provided by the manufacturer (Invitrogen) and 1 µl of tenfold dilutions of the cDNAs synthesized as described above. Thermal cycling parameters were as follows: 94 °C for 2 min for initial denaturation; 40 cycles of 94 °C for 10 s; and 58 °C for 15 s; 72 °C for 20 s for amplification products by PCR and 5 min at 72 °C for final elongation. Denaturation curves for each PCR reaction were determined by measuring the decrease in fluorescence with increasing temperature from 65 to 98 °C each 0.2 °C. The *UBQ3* gene (Solyc01g056940) was used as reference (Hoffman et al., 1991; Hackel et al., 2005). Relative gene expression was calculated using a modified version of the comparative 2^−ΔΔ^CT method (Pfaffl, 2001). The efficiencies were calculated as described by Liu and Saint (2002), and the error propagation according to Hellemans et al. (2007) for 3 to 5 biological replicated used. The oligonucleotide pairs used are in Table S1.

### Heterologous expression and purification of SlNADP-MEs

Plasmids encoding for SlNADP-ME1, -2, -3 and mature -4a and -4b were produced using the BioCat commercial gene synthesis service (Heidelberg, Germany). The coding sequences were synthesized, cloned into pET16b via NdeI/XhoI restriction sites and sequenced for verification.

All expression constructs were transformed into chemically competent Escherichia coli ArcticExpress (DE3) cells (Agilent Technologies, Santa Clara, USA). Transformed cells were grown on LB agar plates containing antibiotics according to the manufacturer’s instruction and 100 µg/ml ampicillin (pET16b selection agent). For the expression of His-tagged SlNADP-ME1, -2, -3 and mature -4a and -4b, LB-cultures containing 20 µg/ml gentamycin and 100 µg/ml ampicillin were grown overnight at 37 °C. Overnight cultures were used to inoculate 400 ml (SlNADP-ME1, -2, and mature -4a and -4b, denoted as protocol 1, P1) and 500 ml (SlNADP-ME3, denoted as protocol 2, P2) of LB-medium. For P1, liquid cell cultures were grown at 30°C and after reaching an OD600 between 0.4-0.6, cultures were cooled down to 11°C. At 11°C, protein expression was induced with 1 mM isopropyl-ß-d-thiogalactopyranoside (IPTG) and cells were incubated under the same conditions for 24 h. For P2, liquid cell cultures were grown at 37°C and after reaching an OD600 between 0.4-0.6, cultures were cooled down to 19°C. At 19°C protein expression was induced with 1 mM IPTG and cells were incubated for 24 h under the same conditions. After incubation, cells were harvested by centrifugation for 10 min at 4°C and 6000 g and stored for further use at -20°C.

### Protein purification

His-tagged proteins were purified using gravity-flow immobilized metal ion chromatography on nickel-nitrilotriacetic acid agarose (Ni-NTA Agarose). Frozen cells were thawed on ice and resuspended in 20 mM Tris-HCl (pH 8.0) containing 500 mM NaCl, 5 mM Imidazole, 2 mM phenylmethanesulfonyl fluoride and a spatula-tip amount of lysozyme. Resuspended cells were sonicated for 8 min (30 s intervals with 30 s pause in between) and centrifuged for 15 min at 15000 g and 4 °C. The supernatant was loaded onto a Ni-NTA agarose (Qiagen, Hilden, Germany) column that was previously equilibrated with Buffer 1 (20 mM Tris-HCl (pH 8.0), 500 mM NaCl, 5 mM Imidazole). In the subsequent steps, the column was washed with Buffer 1 containing increasing imidazole concentrations (5, 40, 80 mM). Before the last washing step, the column was washed with Buffer 2 (20 mM Tris-HCl (pH 8.0) and 2 M Urea) to remove Cpn60. Proteins were eluted with 2 ml elution buffer (20 mM Tris-HCl (pH 8.0), 500 mM NaCl and 500 mM Imidazole) and concentrated using Amicon Ultra 0.5 ml Centrifugal Filters (50 kDa molecular weight cut-off) (Merck, Darmstadt, Germany) according to the instructions of the manufacturer. Concentrated proteins were stored in storage buffer (20 mM Tris-HCl (pH 7.5), 5 mM MgCl2) at 4 °C overnight and in storage buffer with 20 % glycerol at -80 °C for the long term.

### Kinetic Measurements

NADP-ME activity measurements were conducted using a Tecan Spark plate reader (Tecan Group, Männedorf, Switzerland) and enzyme activity (oxidative decarboxylation of L-malate) was determined spectrophotometrically by measuring the formation of NADPH at 340 nm and 25 °C. The Michaelis constants (Km) for malate and NADP were determined by varying the concentrations of malate and NADP around the Km values of the studied protein while keeping the other components at saturating concentrations. A standard reaction mixture contained 50 mM Tris-HCl (pH 7.5), 10 mM MgCl2, 200 ng of protein and either 0.5 mM NADP and a varying amount of L-malate (malate concentrations varied between 0.01 mM and 40 mM) or 10 mM L-malate and a varying amount of NADP (NADP concentrations varied between 0.005 and 0.35 mM for SlNADP-ME1, -3 and 4b, between 0.01 and 0.4 mM for SlNADP-ME2 and between 0.002-0.125 mM for SlNADP-ME1). To measure protein activity, 192 µl of assay solution was prepared in a Greiner 96 well Flat Transparent Plate (Thermo Fisher Scientific, Waltham, Massachusetts) and after a baseline measurement, 8 µl of increasing L-malate or NADP concentrations were added to the different wells to a final assay volume of 200 µl. The formation of NADPH was measured at 340 nm and 25 °C for 25 minutes. All kinetic measurements were performed the next day after purification, with proteins stored at 4 °C overnight. Activity measurements were performed at least three times with independently expressed and purified proteins, with each measurement consisting of a triplicate for each concentration.

### Data analysis

Plate reader data showed the increase of absorbance of NADPH at varying concentrations of malate or NADP. Kinetic parameters were calculated at least three times with independently expressed and purified proteins, with each calculation consisting of a triplicate for each concentration. The linear regression and subsequently the highest slope of the absorbance increase were determined, and triplicates were averaged. From the average absorbance, the specific reaction velocity V0 (mmol*s-1*L-1) was determined using the extinction coefficient (ε) of 6.22 mM-1 cm-1 at 340 nm for NADPH and the formula:

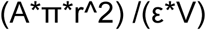

A = absorbance

π = mathematical constant pi

r = radius of the well (0.35 cm)

ε = molar absorption coefficient of NADPH (6.22 mM-1 cm-1 at 340 nm) V = volume of the reaction mixture (200 µL)

The specific reaction velocities at different malate or NADP concentrations were adjusted to non-linear regression in GraphPad Prism 8 (v8.3.0) to obtain the Michaelis-Menten kinetics (Vmax(mmol*s-1*L-1), K_m_ (mM)). Complexation of malate and NADP by Mg^2+^ was accounted for as described in Grover et al. (1981). The catalytic rate (k_cat_ (s-1)) was calculated by dividing Vmax through the amount of enzyme (mol) used in the assay and the catalytic efficiency (k_cat_/K_m_(s^−1^mM^−1^)) was determined by dividing K_m_ by k_cat_.

### Retrieval of promoter regions and analysis of cis-regulatory elements

Promoter sequences (1500 bp upstream and 100 pd downstream of translation start site) of each NADP(P)-ME gene of S. lycopersicum were retrieved www.phytozome.net. The tool PlantCare (http://bioinformatics.psb.ugent.be/webtools/plantcare/html/) were used for scanning of cis-elements present in promoter regions (Supplementary Data 2).

### Phylogenetic and molecular evolutionary Supplementary Data analyses

Sequence retrieval for phylogenetic analysis, NADP-ME coding sequences of plant species with entire genome information were extracted from primary gene models www.phytozome.net. BLASTP with the BLOSUM62 as default scoring matrix and a minimal e-value of 0.0001 was implemented to obtain homologs using SlNADP-ME2 (Solyc05g050120) as query. All sequences were manually checked for correct translation start sites and the presence of conserved amino acid regions found in all NAD(P)-ME (Tronconi et al., 2018).

Multiple Sequence Alignments (MSA) of the coding sequences were assembled using MEGA X (v.10.0.5) (Kumar et al., 2018). The sequences were then translated into amino acids and aligned using Muscle (Edgar, 2004) with the gap opening penalty value of -2.9 and without penalizing its extension. Once retranslated into nucleotides, the alignment was manually edited to select the most-reliable positions in the alignment, assisted by Gblocks1 and TrimAl2 programs. Since the different phylogenetic methods consider columns with gaps in different ways, we applied a stringent criterion by eliminating codons with coverage less than 95%. The final multiple sequence alignment (MSA) consisted of 345 coding sequences from 152 eudicots species with 1.640 nucleotides positions corresponding to 546 codons.

Maximum likelihood (ML) and neighbor joining (NJ) analyses on the whole Supplementary Data set were conducted using MEGA X (v.10.0.5). The goodness of fit of each model to the Supplementary Data was measured by the Bayesian information criterion (BIC) and the model with the lowest BIC score was considered the best description for a specific substitution pattern. The initial tree for the ML search was generated automatically by applying the NJ and BIONJ algorithms, and its branch lengths were adjusted to maximize the likelihood of the Supplementary Data set for that tree topology under the selected model of evolution. Heuristic searches were conducted with the initial tree based on the nearest neighbor interchange (NNI) search where the alternative trees differ in one branching pattern. The reliability of interior branches was assessed with 2,000 bootstrap (B) re-samplings. Nodes with MLB or NJB values 50–69% were regarded as weakly supported, 70–84% as moderately supported, and 85–100% as strongly supported (Hillis and Bull, 1993). The files were saved in Newick format (.nwk) containing all the relevant clade support values and branch length information. The trees were displayed using the FigTree v1.4.4 software.

### Synteny analysis

The protein sequences of three Arabidopsis NADP-ME genes (At2g19900, At5g11670, At5g25880 – corresponding to AtNADP-ME1, AtNADP-ME2, and AtNADP-ME3, respectively) and of three tomato NADP-ME copies (Solyc12g044600, Solyc05g050120, Solyc08g066360 – corresponding to SlNADP-ME1, SlNADP -ME2, and SlNADP -ME3, respectively) were retrieved from NCBI. These sequences were used to retrieve NADP-ME homologs across 310 angiosperm genomes (Supplementary Data 1) through a BLAST search (E-value: 1E-3) (Altschul et al., 1990). The identified homologs were further filtered through the annotation of InterPro protein domains to both queries and subjects by using HMMER3 (default parameters; Mistry et al., 2013) and the removal of subjects whose protein domains did not match the ones of the respective queries (custom R script). In parallel, the PEP and BED files of the 310 genomes used to search for NADP-ME genes were used to compute all-vs-all genome synteny, by using the synteny network pipeline developed by Zhao and Schranz (2017). The obtained synteny network was filtered for the NADP-ME genes retrieved by BLAST and HMMER3 search, and subsequently clustered into syntenic communities (again by following the approach developed by Zhao and Schranz (2017).

### Statistical analyses

Statistical analysis for all experiments was performed using Sigma Plot v 15.0 software using a two-way analysis of variance (with α< 0.05) and Holm–Sidak test for multiple comparisons. Analyses were performed on a row data set before logarithmic transformation for better graphical visualization.

## ACKNOWLEDGEMENTS

We thank Professor Veronica Maurino (Molecular Plant Physiology, IZMB, University of Bonn, Germany) for her critical and constructive comments, which have considerably improved the arguments here presented. This work was supported by the National Agency for Scientific and Technological Promotion PICT2018-1064 to MT and CONICET.

## SUPPORTING INFORMATION

Additional Supporting Information may be found in the online version of this article.

Supplementary Data 1: The genomes used for the synteny analysis and copy number of NADP-ME genes across the syntenic communities.

Supplementary Data 2: Name motif, consensus sequence, position and function of *cis*-elements presents in *NAD(P)-ME* promoters.

## SUPPLEMENTARY FIGURES

**Figure S1.**
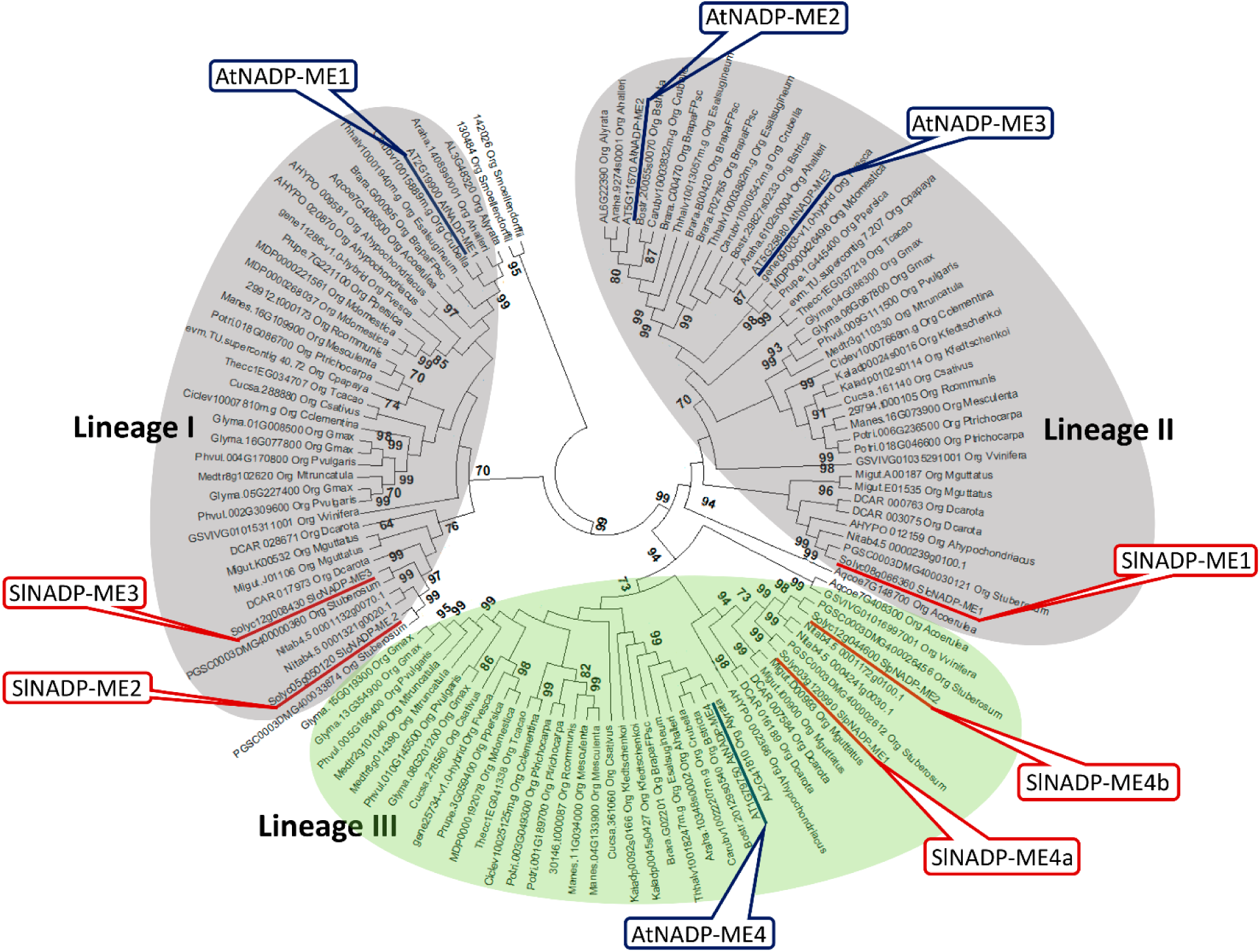
Evolutionary history of NADP-ME genes in eudicot deduced from third position of the codons. Bayesian phylogenetic tree of NADP-ME of eudicot is shown. BPP (Bayesian Posterior Probability) values higher than 70% are given next to the branches. The tree is drawn to scale with branch lengths measured in the number of substitutions per site. The best-fit substitution model was a GRT + G (3.70) model involving 152 nucleotide sequences and a total of 1640 positions in the final Supplementary Dataset. *S. moellendorffi* NADP-ME coding sequences were used as an outgroup.

**Figure S2.**
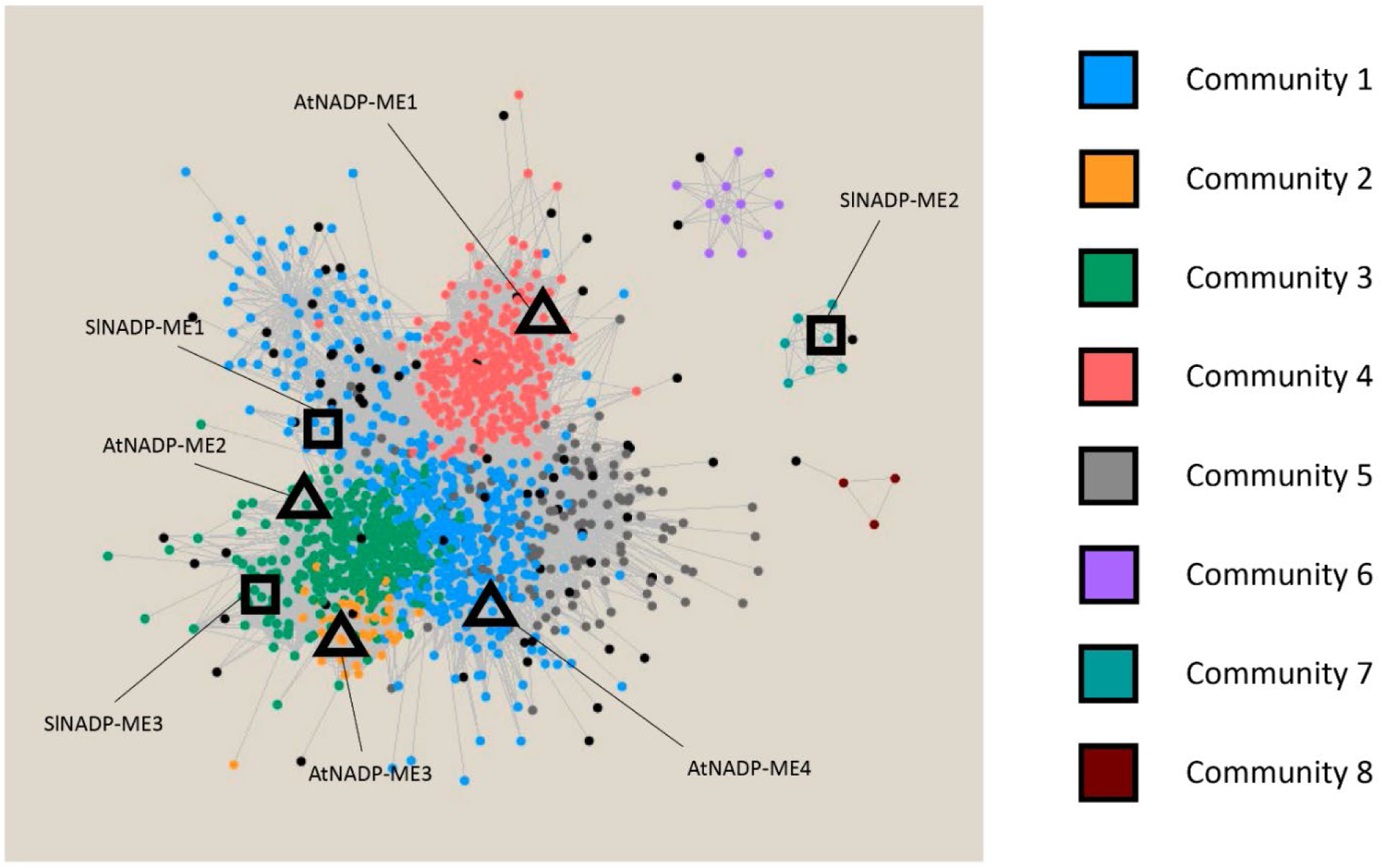
Synteny network of 1400 *NADP-ME* genes across 310 angiosperm genomes. Within the network, nodes are genes and edges represent syntenic connections between genes. Gene nodes are colored based on syntenic communities. Each syntenic community represents an independent genomic configuration, conserved across specific sets of taxa (see supplementary table y for the specific indication of species contained within each syntenic community, and gene copy number across species and syntenic communities). Squares and triangles highlight nodes corresponding to tomato and Arabidopsis *NADP-ME* genes, respectively.

**Figure S3.**
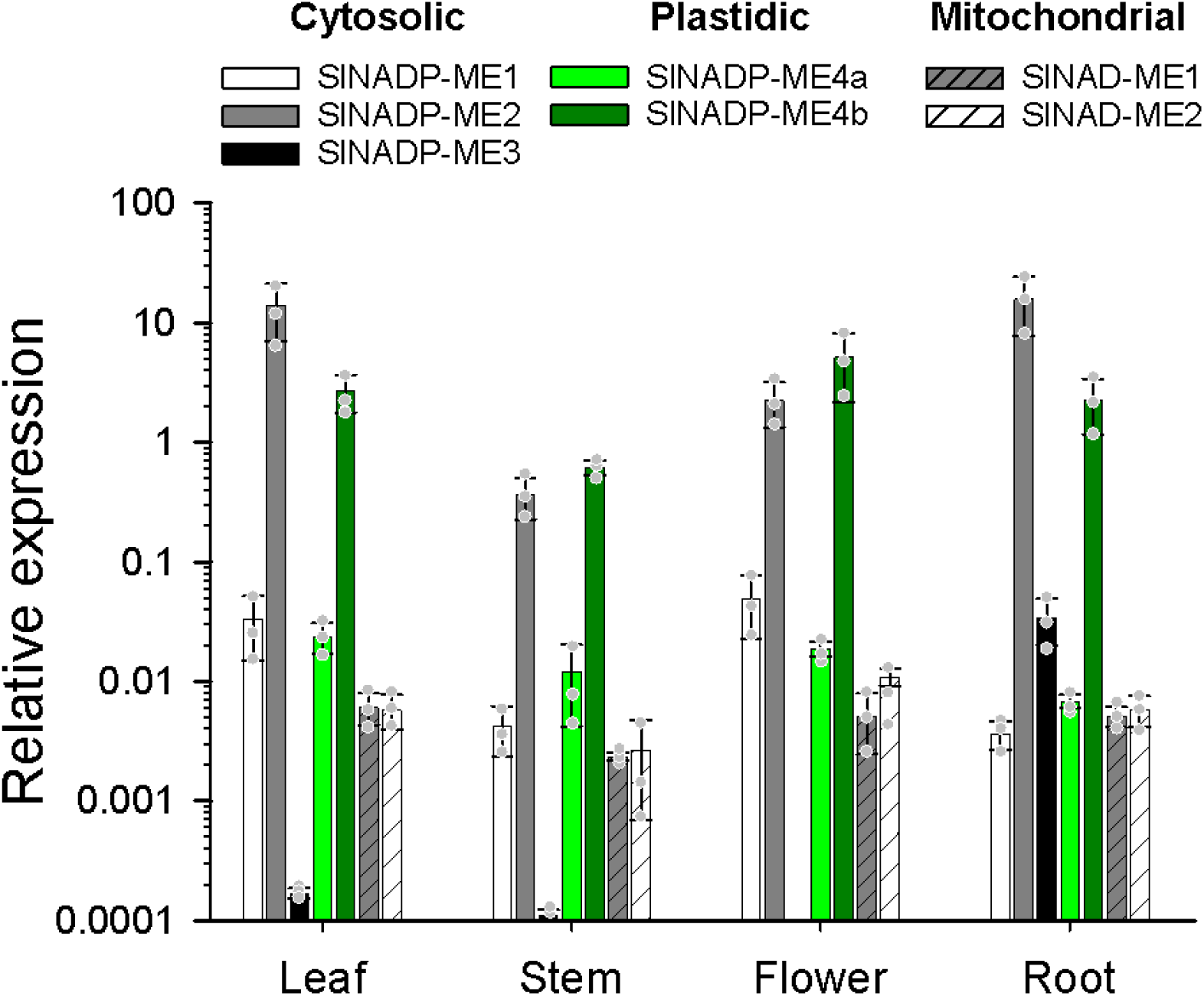
Expression levels of *SlNAD(P)-ME* genes in organs of tomato plants. Transcript abundance was determined by qPCR and the values expressed relative to the abundance of the UBQ3 transcript. Bars represent the mean ± s.d of three independent biological replicates (n=3). Organs from 5-week-old plants were analyzed. The y-axis is shown on a logarithmic scale.

**Figure S4.**
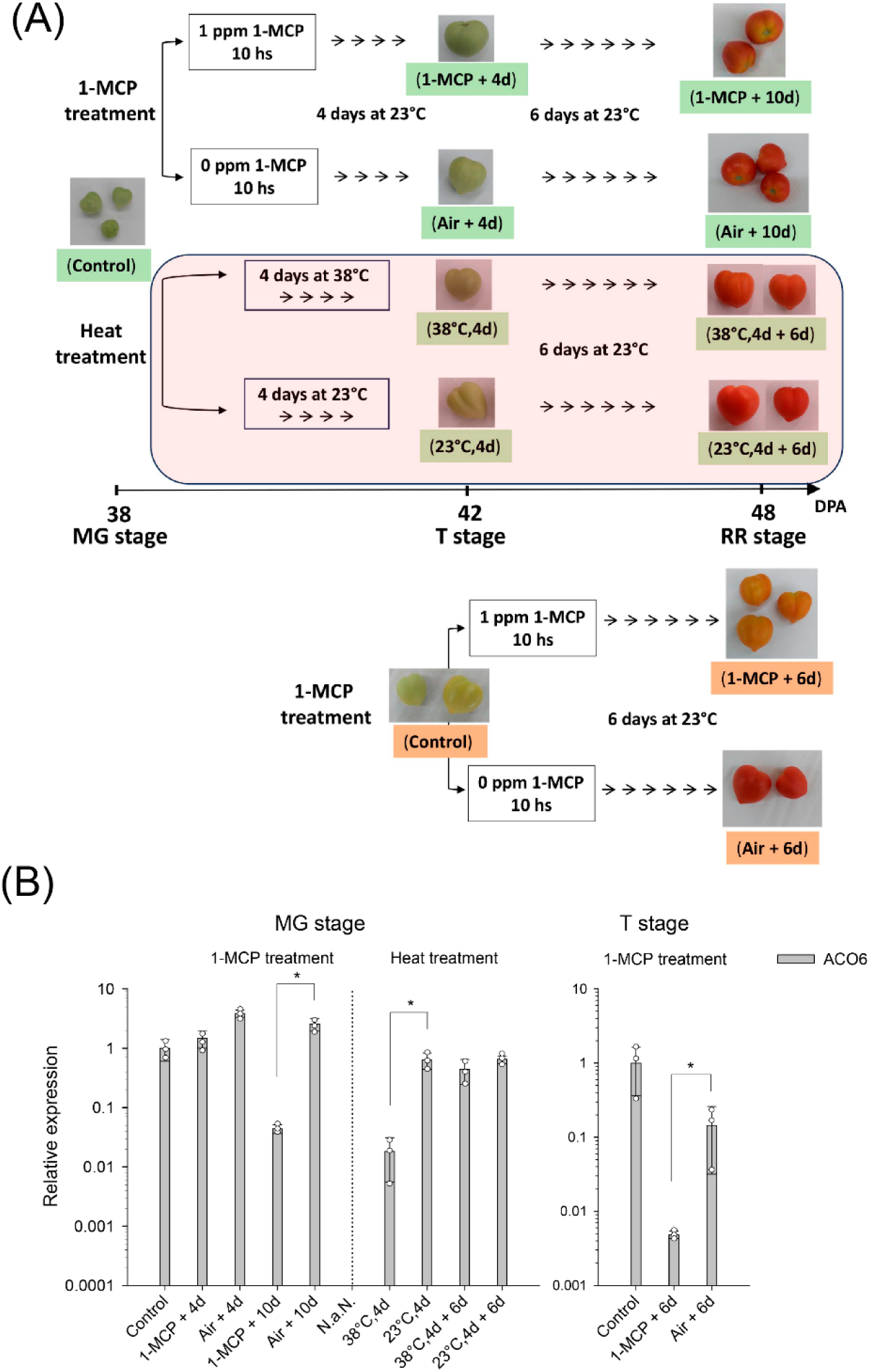
(**A**) Scheme of sample collection and treatment. Mature green and turning fruits were collected from tomato plants at harvest (Control) and treated with 1 ppm 1-MPC for 10 h or incubated at 38°C for 4 days. Sample (pericarps) were taken after 4- and 10-days post-harvest. (**B**) Expression levels of *ACO6* gene in pericarp of fruits 1-MCP-treated or held at 38°C. The levels of transcripts were assessed by qPCR and values expressed relative to the abundance in control. Bars represent the mean ± s.d of three independent biological replicates (n=3). Statistical differences (student’s t test, p<0.05, n=3) between treated and untreated samples are shown as *. The y-axis is shown on a logarithmic scale.

**Figure S5.**
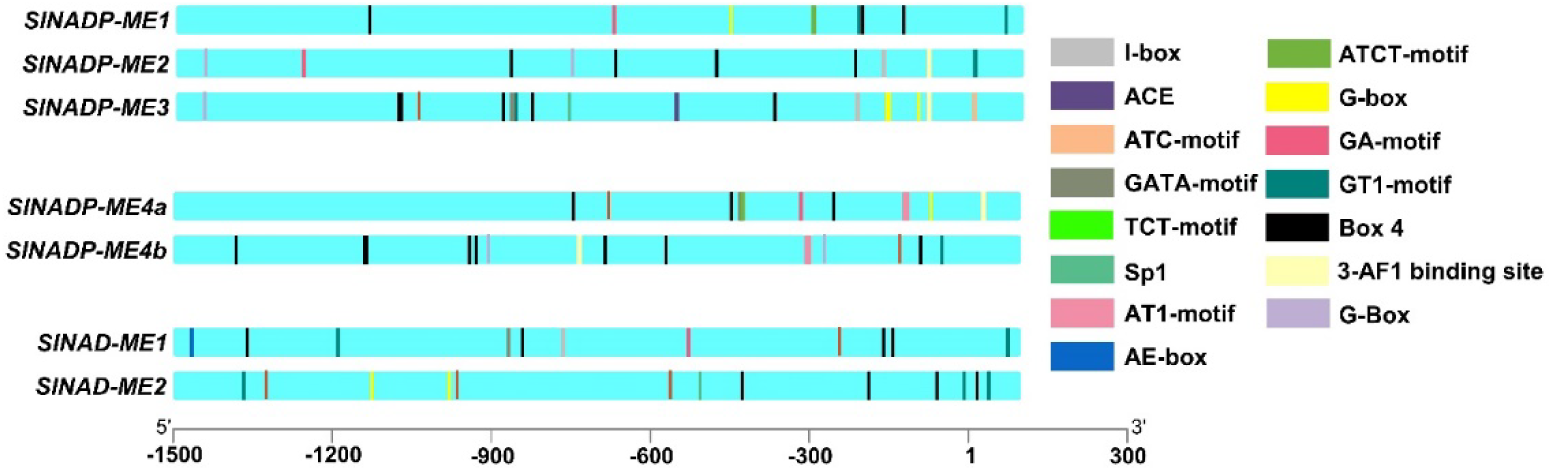
Position of identified Light responsive *cis*-elements in *SlNAD(P)-ME* promoters. TSS (star transcription site) is numbered as 1.

**Figure S6.**
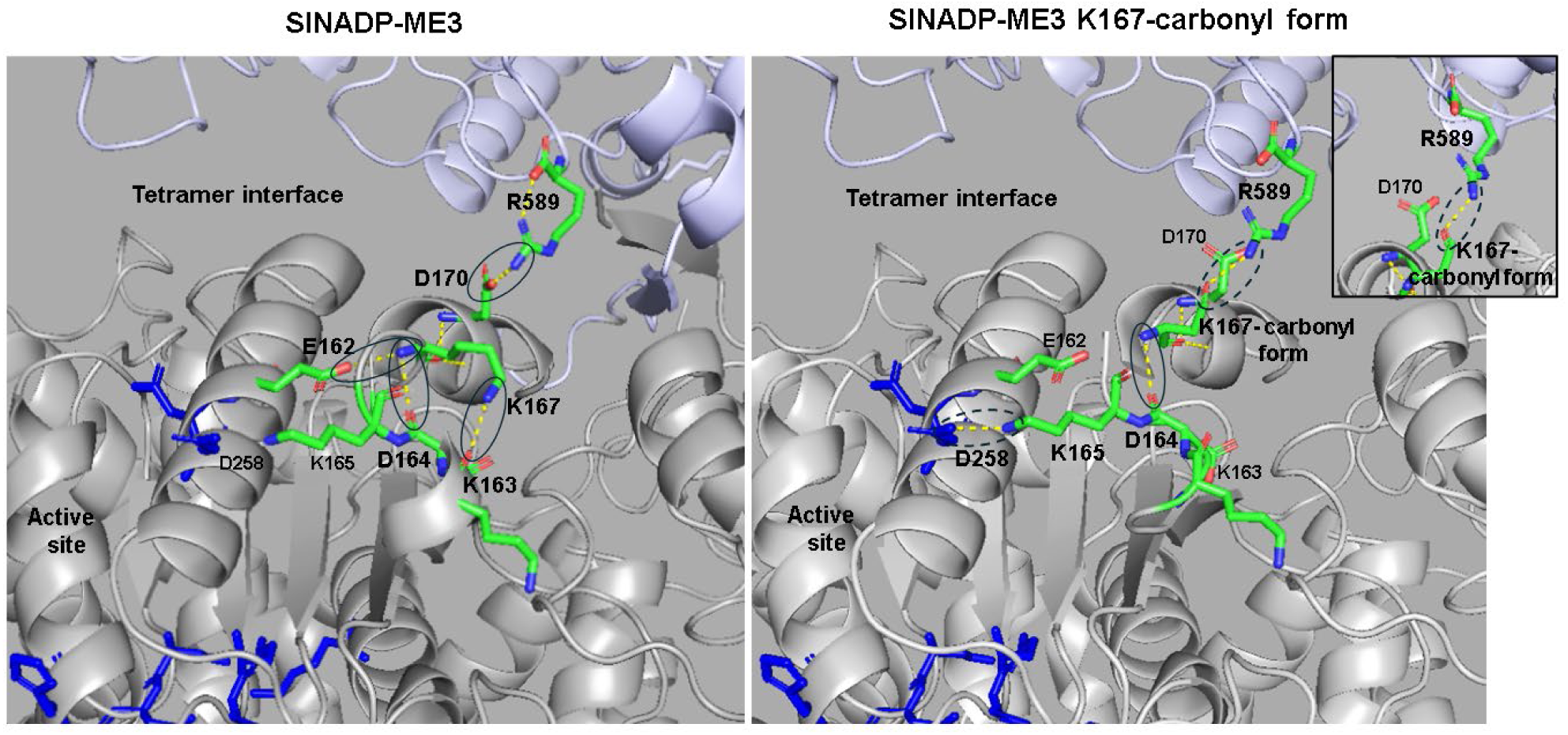
Key structural differences between SlNADP-ME3 and SlNADP-ME3 K167-carbonil form. Comparative modeled structural representation showing significant changes in the tetramer interface by K167-carbonilation of SlNADP-ME3. Interacting monomers are depicted in grey and light blue. Residues associated at the active site are shown in blue. Pairs of interacting residues are enclosed within an oval. Dash line ovals indicate interactions only present in the carbonyl form of SlNADP-ME3. The R589-D170 inter-subunit interaction in non-modified SlNADP-ME3 (left panel) is no longer conserved in the carbonylated form. Instead, R589 interacts with the carbonyl group of K167-semialdheide (right panel). The inset in the right panel shows the interaction R589-K167-carbonylated from a different angle. The models were generated using the protein structure homology-modelling SWISS-MODEL server (swissmodel.expasy.org) and the resolved structure of C4-NADP-ME from maize as templated (PDB ID: 5OU5, Alvarez et. Al., 2019).

**Table S1.**
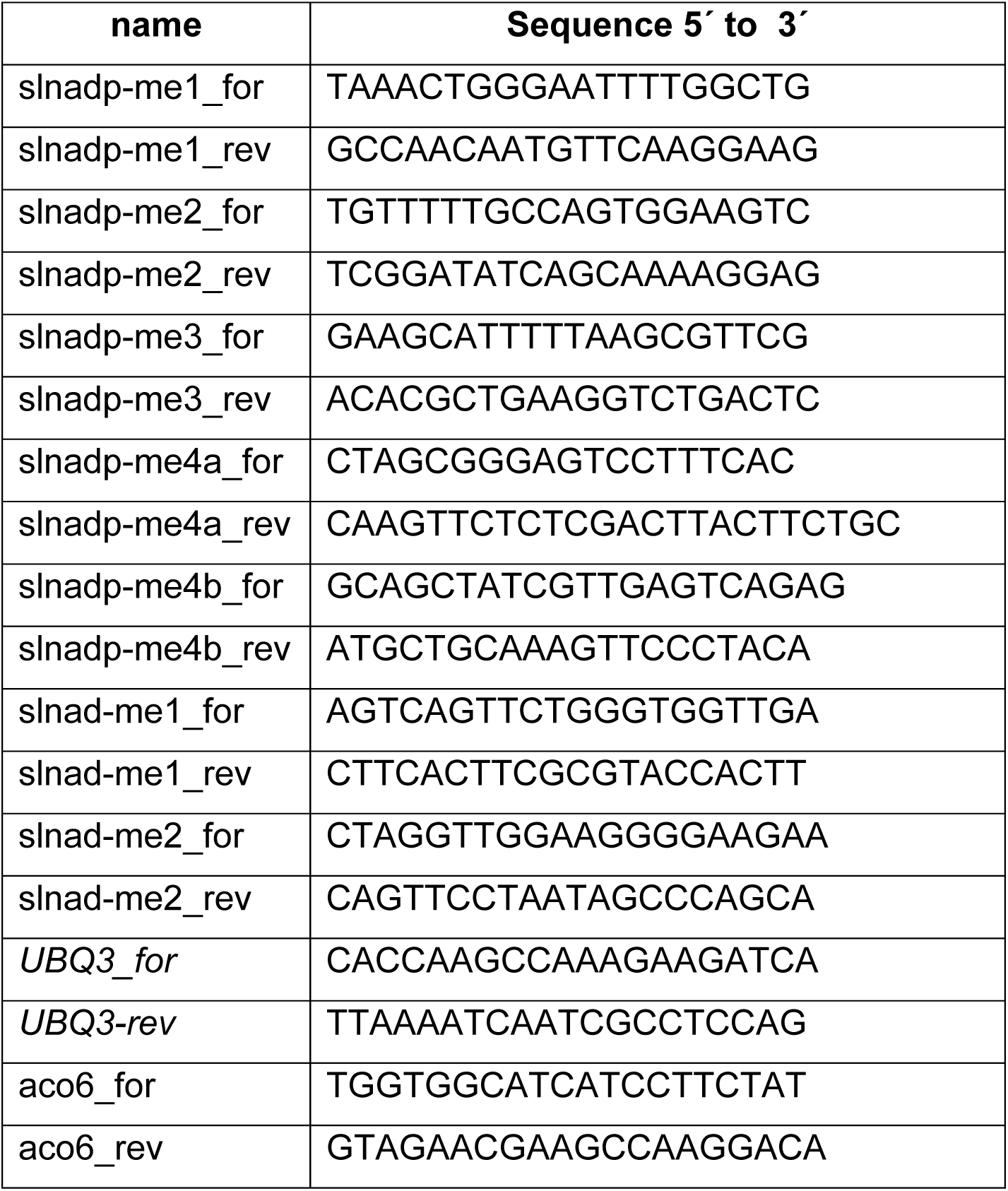
Nucleotides used in this work for RT-qPCR.

